# The Role of Ryanodine Receptor 2 in Drug-Associated Learning

**DOI:** 10.1101/2023.10.03.560743

**Authors:** Kara R. Barber, Velia S. Vizcarra, Ashlyn Zilch, Lisa Majuta, Cody C. Diezel, Oliver P. Culver, Brandon W. Hughes, Makoto Taniguchi, John M. Streicher, Todd W. Vanderah, Arthur C. Riegel

## Abstract

Type-2 ryanodine receptor (RyR2) ion channels facilitate the release of Ca^2+^ from stores and serve an important function in neuroplasticity. The role for RyR2 in hippocampal-dependent learning and memory is well established and chronic hyperphosphorylation of RyR2 (RyR2P) is associated with pathological calcium leakage and cognitive disorders, including Alzheimer’s disease. By comparison, little is known about the role of RyR2 in the ventral medial prefrontal cortex (vmPFC) circuitry important for working memory, decision making, and reward seeking. Here, we evaluated the basal expression and localization of RyR2 and RyR2P in the vmPFC. Next, we employed an operant model of sucrose, cocaine, or morphine self-administration (SA) followed by a (reward-free) recall test, to reengage vmPFC neurons and reactivate reward-seeking and re-evaluated the expression and localization of RyR2 and RyR2P in vmPFC. Under basal conditions, RyR2 was expressed in pyramidal cells but not regularly detected in PV/SST interneurons. On the contrary, RyR2P was rarely observed in PFC somata and was restricted to a different subcompartment of the same neuron - the apical dendrites of layer-5 pyramidal cells. Chronic SA of drug (cocaine or morphine) and nondrug (sucrose) rewards produced comparable increases in RyR2 protein expression. However, recalling either drug reward impaired the usual localization of RyR2P in dendrites and markedly increased its expression in somata immunoreactive for Fos, a marker of highly activated neurons. These effects could not be explained by chronic stress or drug withdrawal and instead appeared to require a recall experience associated with prior drug SA. In addition to showing the differential distribution of RyR2/RyR2P and affirming the general role of vmPFC in reward learning, this study provides information on the propensity of addictive drugs to redistribute RyR2P ion channels in a neuronal population engaged in drug-seeking. Hence, focusing on the early impact of addictive drugs on RyR2 function may serve as a promising approach to finding a treatment for substance use disorders.

## Introduction

Ryanodine receptors (RyR) are Ca^2+^ release channels that are expressed from early development[1,2] in highly interconnected membranes of the sarcoplasmic/endoplasmic reticulum[3]. RyR2 is the dominant RyR isoform in the brain, with dense expression in the hippocampus and cortex[4–6]. In the hippocampus, activation of dendritic RyR2 produces regenerative waves of Ca^2+^ release from intracellular stores[7]. These Ca^2+^ waves amplify somatic RyR2 and nuclear Ca2+ signals[8,9] to trigger activity-dependent changes in gene expression that upregulate RyR2 expression[10,11] and transcription factors (e.g., p-CREB, Npas4)[12] important for spatial learning and memory[12,13]. In contrast, decreases in RyR2 expression reduce spatial memory and spine remodeling[9,12,14,15]. In memory tasks, kinase activity is associated with changes in RyR2 channel expression [12,16], and constitutive basal phosphorylation of RyR2 is reported across species[17–20]. However, at a population level, PKA “hyperphosphorylation” of RyR2 serine residue 2808[21,22] is a postulated mechanism for Ca^2+^ leak from intracellular stores in both hippocampal[15] and cortical[23] neurons, as well as stress-induced cognitive dysfunction, including learning deficits[15,24,25].

In the prefrontal cortex (PFC), a region associated with executive function, choice behaviors, and reward-seeking, including addiction, the role of RyR2 is less clear. Human neuroimaging studies show that cocaine- and heroin-paired cues activate the PFC[26,27] and that the magnitude of this activation predicts craving and the risk of relapse[27,28]. In preclinical rodent models, behaviorally relevant cues that predict rewards activate subpopulations of PFC neurons[29–31] believed to store the memory trace of such learned associations[32–34]. These neuronal ensembles can be identified using immediate early gene indicators such as Fos, whose expression increases following strong neural activation[35–37]. Relative to nondrug reinforcers (i.e., food or sucrose), addictive drugs produce robust, long-lasting neural changes that result in stronger, more persistent behavioral responses for drugs[38,39]. Interestingly, cocaine increases RyR2 expression in cortical neuron cultures[40]. Repeated intraperitoneal (IP) injections of nicotine[13] or methamphetamine[41] also increase RyR2 levels in the frontal cortex, and nonselective inhibition of RyR by intracerebroventricular (ICV) application of dantrolene is reported to block methamphetamine CPP[41]. These studies suggest the functional involvement of RyR2 in drug-induced neuroplasticity, although any specific role in drug-seeking or hyperphosphorylation has not been investigated.

Here, we hypothesized that increased RyR2 hyperphosphorylation could promote addictive behaviors independent of the drug’s other effects on total RyR2 protein expression. To investigate this idea, we mapped the basal expression and localization of total RyR2 (TRyR2) and RyR2P in the vmPFC. Then, using a rat model of chronic self-administration (SA), we tested the animals’ recall behavior while seeking a drug reward (cocaine or morphine) and compared the TRyR2/RyR2P responses to that of a nondrug/natural (sucrose) reward. Although recalling both types of rewards enhanced TRyR2 expression, our results revealed that only addictive drugs were associated with a redistribution of RyR2P from active dendrites to cell bodies colocalized with Fos, a marker of highly activated neurons[30,42]. These adaptations were not observed with stress or noncontingent drug exposure and were independent of changes in TRyR2 transcription. Our concluding observations shed light on a learning-related, drug-specific posttranslational modification of RyR2 in a discrete population of excitable neurons in a brain region classically associated with drug-seeking.

## Results

We combined operant behavior paradigms involving self-administration (SA) of drug and nondrug rewards, along with immunohistochemistry, Western blot analysis, and RNAScope techniques to address the following questions: (1) What enduring modifications occur in the ventral medial PFC (vmPFC) after the acquisition and recall of a learned event? (2) Do RyR2 responses differ between drug and nondrug rewards? (3) Does the recall process of such learned events induce changes in expression patterns that potentially enhance vmPFC activity? Using these methods, we examined whether RyR2 expression in vmPFC differed after nondrug and drug rewards, and whether recalling the memory of these previous experiences impacted vmPFC function.

### Basal RyR2 expression and phosphorylation in vmPFC

Using vmPFC sections from naive adult male rats, we performed immunohistochemical staining with RyR2-specific antibodies[43]. Confocal analysis indicated robust somatodendritic total RyR2 (TRyR2) expression (**Figure 1A**) in the soma and apical dendrites of layers 2-3 (L2-3) as well as in the pyramidal neurons of L5 PFC. Using calcium/calmodulin-dependent protein kinase II (CaMKII) as a marker of cortical glutamatergic pyramidal projection neurons[44–46], we performed double-label fluorescent immunohistochemistry studies to identify CaMKII or co-labeled TRyR2+CaMKII-immunoreactive neurons (**Figure 1B**). Quantification of the two markers indicated significant colocalization in the vmPFC (**Figure 1C**). In contrast, TRyR2 showed little overlap with parvalbumin (PV)[47,48] or somatostatin (SST)[49] inhibitory neurons (**Figure 1-figure supplement 1 and Figure 1C**) (one-way ANOVA, F_(2,23)_=56.88, p<0.0001, followed by Tukey’s multiple comparison test: CaMKII vs. PV, p<0.0001; CaMKII vs. SST, p<0.0001; PV vs. SST, p=0.9484). Therefore, as in other regions[9,50], excitatory pyramidal cells in vmPFC possessed a high basal expression of TRyR2.

**Figure 1.**
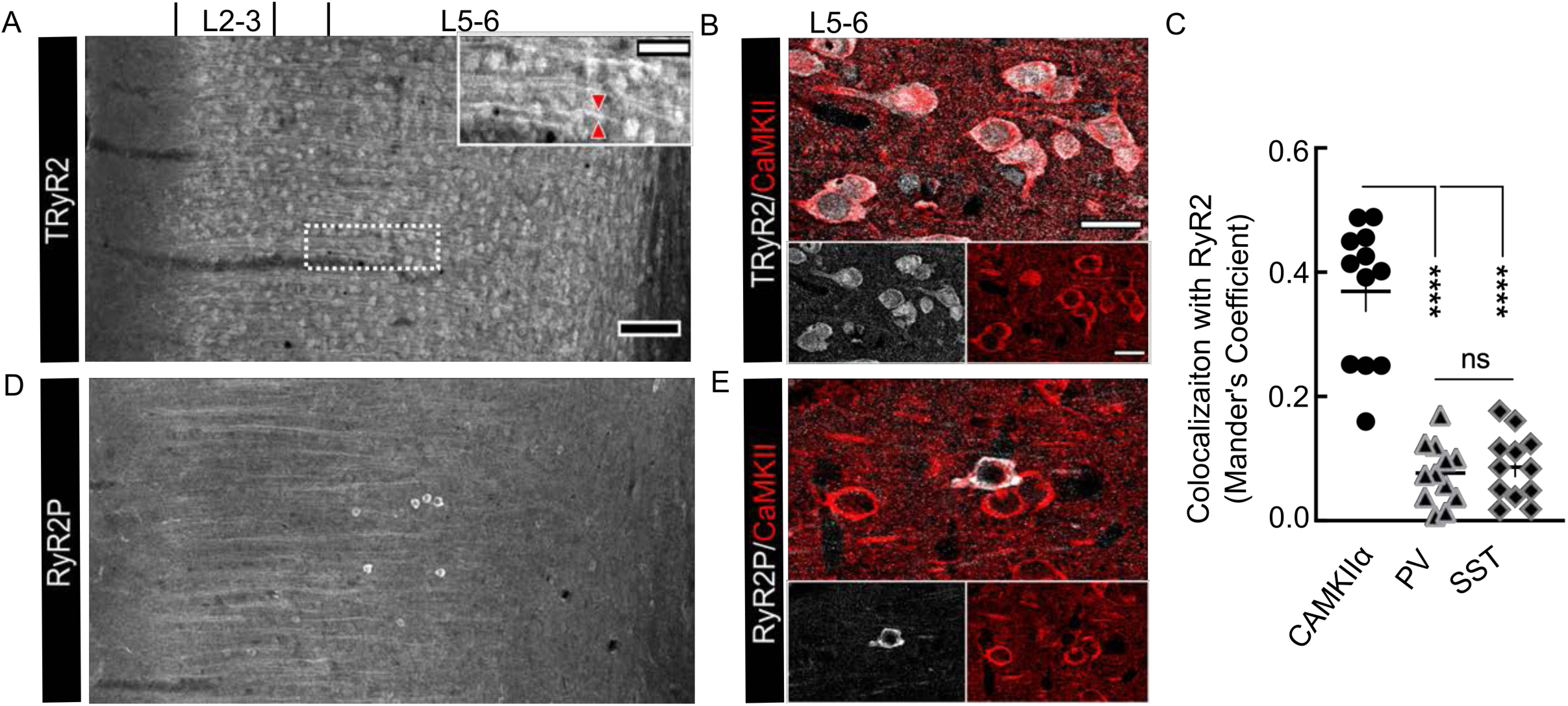
TRyR2 and RyR2P are distributed in a somatodendritic pattern in excitatory PFC neurons. Confocal photomicrographs of a stack of serial vmPFC sections showing (**A**) high expression of TRyR2 (*white*) in dendrites (see *inset red arrowheads;* scale bar = 50 μm) and somata of L2-3 and L5-6 pyramidal cells (scale bar 100 μm) that (**B**) colocalized with CaMKII (*red*), a marker of excitatory pyramidal neurons in the PFC (scale bar = 20 μm). (**C**) Quantification of Manders’ coefficient showing the percent overlap of TRyR2 and CAMKII (one-way ANOVA, F(2,23)=56.88, p<0.0001, followed by Tukey’s multiple comparison Test; CaMKII vs. PV, p<0.0001; CaMKII vs. SST, p<0.0001; PV vs. SST, p=0.9484). (**D**) In contrast, vmPFC RyR2P (*white*) expressed predominantly in apical dendrites that travel from L5 to L2-3 and (**E**) seldom colocalized with CaMKII (*red*).

To test for phosphorylation of RyR2 (RyR2P), we first replicated a previous study evaluating the specificity of a phospho-antibody directed against RyR2P[51,52] **(Figure 1-figure supplement 2**). We transfected human embryonic kidney cells with wild-type RyR2 (WTS2808) plasmids or a mutant RyR2 S2808 converted to an alanine (MTS-A2808) that prevents phosphorylation. Then we used double-label fluorescence immunohistochemistry to evaluate RyR or co-labeled RyR2P-immunoreactive cells, with or without forskolin (10 µM; 1 hr), an activator of adenyl cyclase that increases PKA activity[53]. Confocal analysis indicated that compared to similarly treated WTS2808 cells, mutant S-A2808 cells expressing RyR showed lower RyR2P fluorescence under basal conditions (∼4-fold; **Figure 1 figure supplement 2A,C)** and in response to forskolin (∼6-fold; **Figure 1 figure supplement 2B,C)**, highlighting the usefulness of the antibody to detect RyR2 phosphorylation at S2808.

We then selected brain sections from naïve rats to identify vmPFC sites with high RyR2 phosphorylation activity. We predicted RyR2P expression would parallel the observed somatodendritic staining of TRyR2 in pyramidal L2,3,5 cells **(Figure 1A)**. However, evaluation of confocal photomicrographs across naïve animals (n=3) showed RyR2P staining only in the apical dendrites **(Figure 1D)** of L5 pyramidal cells, an important subcompartment for regenerative electrical events that integrate sensory inputs[54–57]. In rare instances where RyR2P was expressed in the pyramidal cell somata of L5 (**Figure 1 D**), we observed that it co-localized with CAMKII (**Figure 1E**) and not the inhibitory neuron markers SST[49] or PV (**Figure 1 figure supplement 1B**). These data indicated that in vmPFC neurons, RyR2 and constitutively active RyR2P were restricted to different neuronal subcompartments under basal conditions.

### Chronic self-administration and recall testing

Neurons in the mPFC exert adaptive control over learned reward-seeking behaviors[58–61]. To model this in rats, we used the self-administration (SA) model[62] and trained rats to perform sucrose-SA (45 mg; n=23), cocaine-SA (0.5 mg/kg/infusion; n=21) with yoked-saline controls (n=14), or morphine-SA (0.5 mg/kg/infusion; n=15) (**Figure 2A**). During SA training sessions (**Figure 2A**), pressing the active lever resulted in light/tone cues paired with the delivery of the reinforcer, while pressing the inactive lever had no programmed consequence. Yoked-saline rats were matched with cocaine-SA rats to receive the same conditions but passive infusions of saline instead of cocaine (**Figure 2B1**). As an additional control, a cohort of cocaine-SA and (control) sucrose-SA rats received either a withdrawal or active drug-taking test on their last session day (**Figure 2A**). Analysis of the resulting line graphs of chronic pressing activity (**Figures 2B2-B4)** indicated that rats readily learned to distinguish between active and inactive levers during the 14 sessions and actively sought the reward (mixed-effects Two-way ANOVA, p<0.0001; sucrose F_(2,66)_ = 68.67; cocaine F_(2,60)_ = 116.9: morphine F_(2,42)_=37.18). Behavioral responding remained stable over time, as previously reported by us[62,63] and others[64–67].

**Figure 2.**
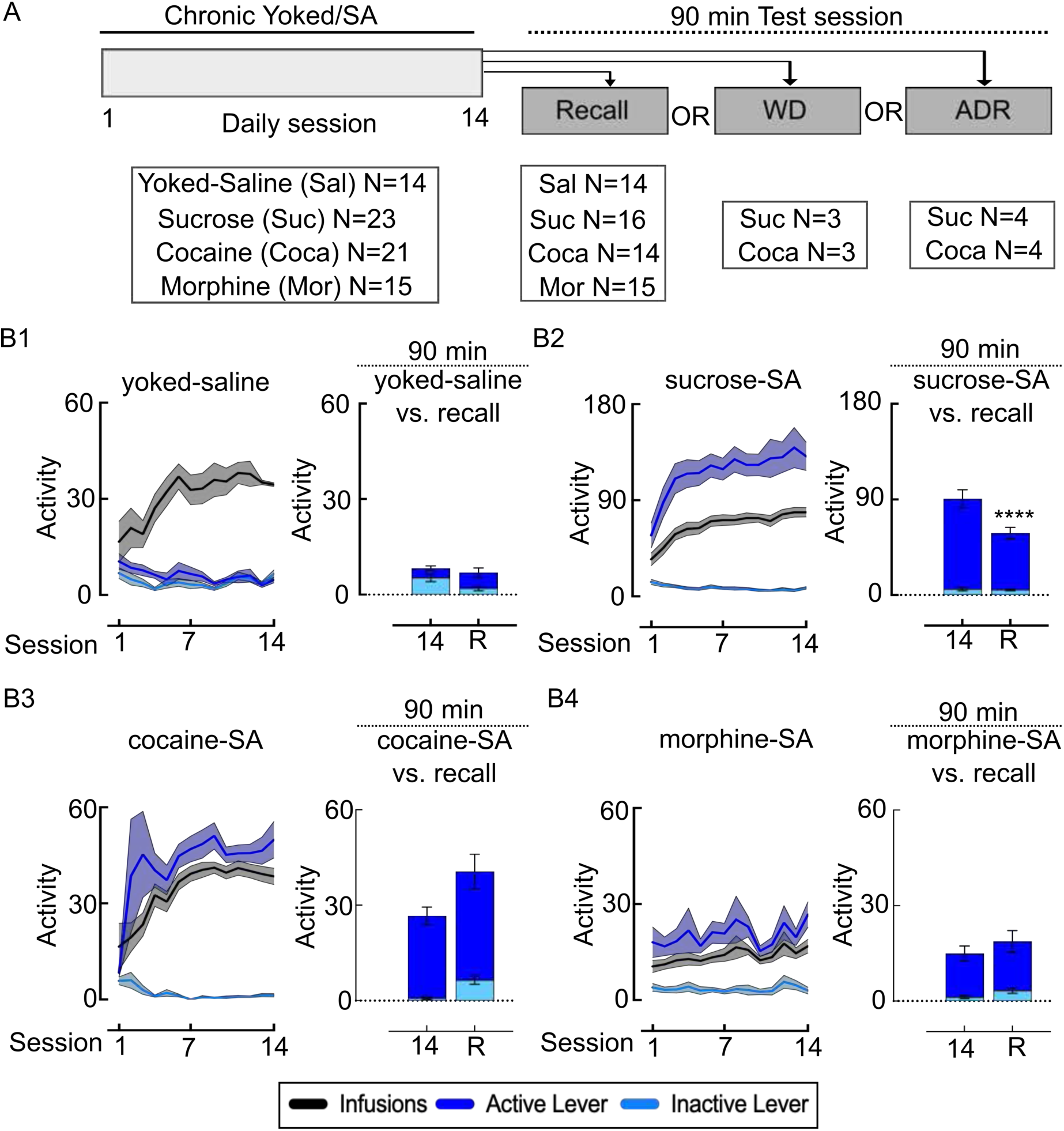
Experimental design for self-administration and recall testing. **(A)** Rats learned to self-administer (SA) a reward (cocaine, morphine, or sucrose) during daily, repeated (14d) sessions. Delivery of cocaine (0.5 mg/kg, i.v.; 2hr/day), yoked-saline (2hr/day), morphine (0.5 mg/kg, i.v; 3hr/day), or sucrose (45 mg pellets; 3hr/day) occurred in an FR1 reinforcement schedule in conjunction with discrete cues (light and tone). On day 15, rats experienced a 90-minute Test session to evaluate: Recall (without reward or discrete cues), Withdrawal (WD) (without reward in homecage), or Active delivery of reinforcer (ADR) (with reward and discrete cues). At 90 min after the Test session, rats received either cardiac perfusion with PBS and 4% PFA (for IF or RNAscope) or rapid decapitation (for IP studies). (**B1-B4**) Line plots summarizing lever pressing activity and infusions in (**B1**) yoked-saline, (**B2**) sucrose-SA, (**B3**) cocaine-SA, or (**B4**) morphine-SA rats. SA rats discriminated between an inactive lever (*light blue*) with no reinforcement or an active lever (*dark blue*) that delivered a reward reinforcement (infusions or pellets; *black*) (mixed-effects Two-way ANOVA, p<0.0001; sucrose = F(2,45)=57.49, cocaine = F(2,41)=36.97, morphine = F(2,39)=34.72). Relative to the first 90 min of SA responding on day 14, recall (*R*) testing (day 15) decreased inactive lever responding in yoked-saline rats (**B1;** day: F_(1,26)_=0.7948, p=0.3808; lever: F_(1,26)_=0.06452, p=0.8015; day X lever, F_(1,26)_=11.73, p=0.0021) and decreased active lever responding in the nondrug (sucrose) rats (**B2;** day: F_(1,30)_=0.19.79, p=0.0001; lever: F_(1,30)_=101.0, p<0.0001; day X lever: F_(1,30)_=18.00, p=0.0002). Relative to day 14, active lever responding during recall was similar or increased in cocaine rats (**B3;** day F_(1,26)_=6.034, p=0.0210; lever F_(1,26)_=59.94, p<0.0001; day X lever F_(1,26)_=0.2056, p=0.6540) and morphine rats (**B4;** day F_(1,14)_=0.7158, p=0.4118; lever F_(1,14)_=41.98, p<0.0001; day X lever F_(1,14)_=0.0029, p=0.9580). Error windows and bars represent S.E.M.

As a test of seeking behavior without the confounding effect of the active drug, some rats underwent a recall test[65] on day 15, 24 hours after their last SA session (**Figure 2A**). Rats were placed in the same SA boxes as training but without reinforcers or cues. As in previous studies[65], we reasoned that for rats trained with various rewards, the predominant memories recalled during the 90-minute test session would be memories associated with sucrose, cocaine, or morphine-SA. In control and nondrug rewards, we compared behavioral responses during recall with the first 90 min of their respective preceding SA session (day 14) using a two-way ANOVA. Recall testing (*R*) produced no significant change in active lever presses and slightly reduced inactive lever responding in yoked-saline rats (**Figure 2B1;** day: F_(1,26)_=0.7948, p=0.3808; lever: F_(1,26)_=0.06452, p=0.8015; day X lever, F_(1,26)_=11.73, p=0.0021). In nondrug (sucrose) rats, recall testing (*R*) produced the expected reduction[68] in active lever responding (p<0.0001) relative to day 14 SA (**Figure 2B2;** day: F_(1,30)_=0.19.79, p=0.0001; lever: F_(1,30)_=101.0, p<0.0001; day X lever: F_(1,30)_=18.00, p=0.0002). In contrast, active lever responding during the recall test (*R*) was similar or increased in the drug reward groups compared to day 14 SA (**Figure 2B3,4),** as previously reported[65] (cocaine: day F_(1,26)_=6.034, p=0.0210; lever F_(1,26)_=59.94, p<0.0001; day X lever F_(1,26)_=0.2056, p=0.6540; morphine: day F_(1,14)_=0.7158, p=0.4118; lever F_(1,14)_=41.98, p<0.0001; day X lever F_(1,14)_=0.0029, p=0.9580). Therefore, the seeking behavior with cocaine or morphine recall was stronger than that for a nondrug reward.

### Recall-dependent localization of RyR2P

In rats, recalling a previous reward experience activates vmPFC neurons[69], but it remains unknown whether nondrug or drug rewards differentially alter RyR2P localization. Therefore, we compared behavioral responses and RyR2P expression in the above yoked-saline, sucrose-SA, and cocaine-SA rats. We incorporated a fluorescent double-label of NeuN with RyR2P immunohistochemistry and at low magnification, aligned hemispheres (bregma = +3.00 - +3.72 mm) per group according to cell layers (1-6) (**Figure 3A**)[70]. Then, using the same confocal settings, we compared the RyR2P immunofluorescence at high magnification between the treatment groups.

**Figure 3.**
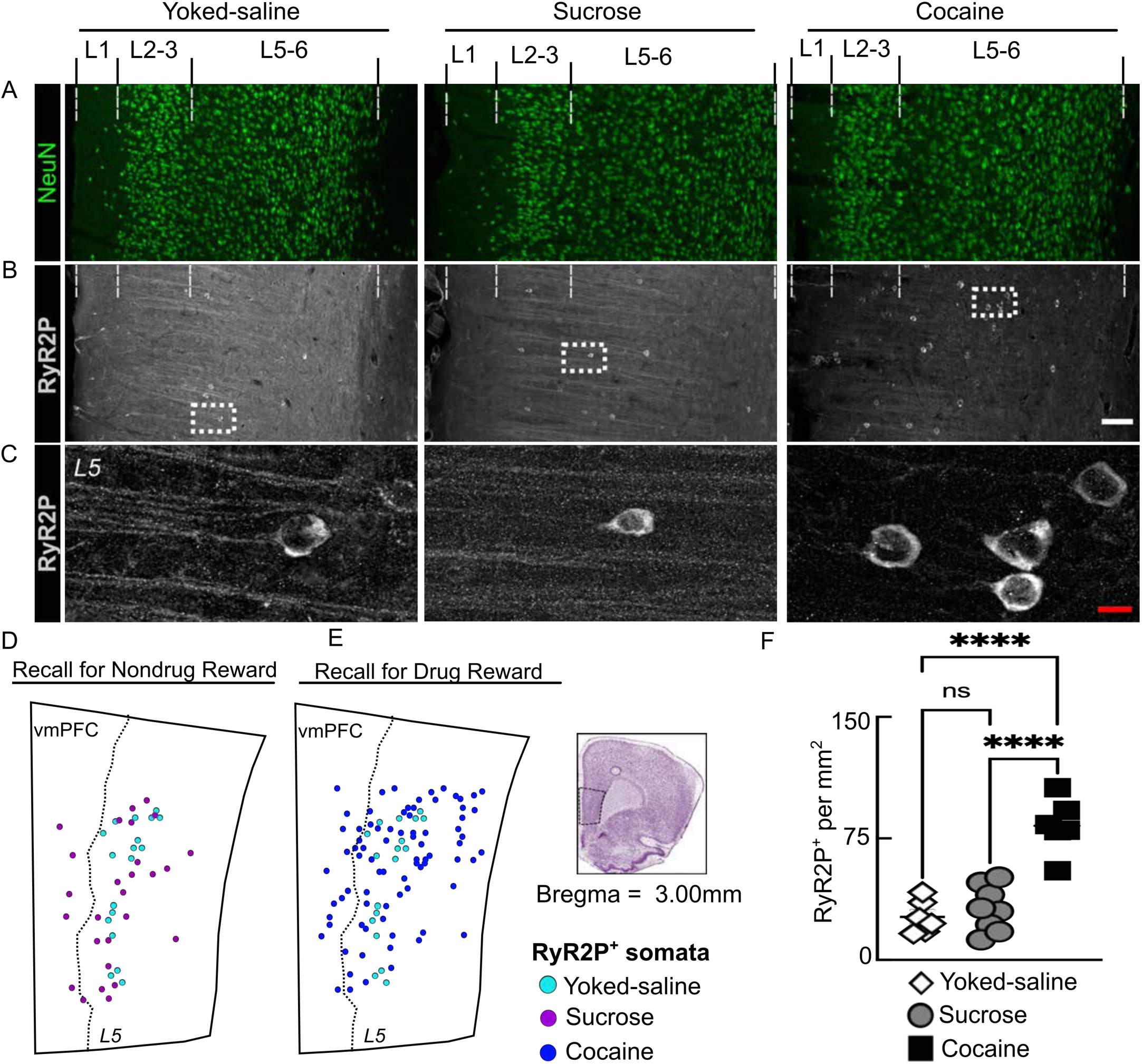
Spatial localization of RyR2P in the vmPFC following recall of natural and drug rewards. (**A-C**) Confocal photomicrographs of vmPFC sections from yoked-saline, sucrose, and cocaine recall rats. (**A-B**) Alignment of 10x images showing L1-6 cells stained with antibodies against NeuN (*green*; **A**) and RyR2P (*white*; **B,C**). (**C**) 60x images of the insets shown in (**B**). Note, the changes in RyR2P expression in apical dendrites (yoked-saline and sucrose) and soma (cocaine). (**D-E**) Cumulative map of RyR2P positive somata in the vmPFC comparing (**D,E**) yoked-saline controls (*light blue dots*) to (**D**) sucrose (*magenta dots*) (**E**)cocaine (*dark blue dots*). The vertical dotted line represents the demarcation of L2-4 (on the *left*) and L5-6 (on the *right*). Scale bar: white = 100 μm; red = 10 μm. (**F**) Summary graph showing the quantification of RyR2P positive neurons per mm2 (One-way ANOVA (F_(2,17)_=30.79, p<0.0001) Error bars represent S.E.M. and Tukey post hoc (ns = not significant, ****=p≤0.0001). N=6-8 rats per experimental group with 4 technical replicates per rat.

Although recall behavior in sucrose rats was elevated over yoked-saline rats (**Figure 2 B1,B2**), we noted similar RyR2P localization in both groups on photomicrographs (**Figure 3 B,C**). In fact, localization in yoked-saline and sucrose recall tissue was not noticeably different from that in the naïve rats (**Figure 1D**). All three groups expressed RyR2P in the apical dendrites (**Figure 3 B,C**). A cumulative map of RyR2P localization indicated similar somatic expression in L5-6 pyramidal cells in sucrose recall tissue (**Figure 3D**, *magenta*) and yoked-saline tissue (**Figure 3 D,E,** *light blue*). That is, recall of a nondrug reward sufficient to activate vmPFC neurons did not noticeably increase RyR2P localization.

To determine whether the type of reward (drug versus nondrug) impacted RyR2P expression, we selected tissue from control (yoked-saline and sucrose-SA) and cocaine-SA rats that had increased responding during recall behavior (**Figure 2 B2-B3**). Comparing these images, we made three principal observations. First, photomicrographs of cocaine-SA recall tissue showed more frequent (**Figure 3 B**, *low magnification*) and more robust (**Figure 3 C,** *high magnification*) somatic RyR2P staining. Second, cumulative mapping (**Figure 3 E,** *dark blue*) linked the recall of cocaine with the expansive expression of RyR2P in L2-3 and L5. Third, in the cocaine recall group, the location of RyR2P changed (**Figure 3C**). That is, RyR2P staining in the apical dendrites evident in tissue from naïve rats (**Figure 1 D**), and the control recall (yoked-saline and sucrose) groups (**Figure 3 B,C** *left* and *center*) was absent in the cocaine recall (**Figure 3 B,C** *right*). Moreover, a one-way ANOVA indicated significant differences in RyR2P somata count between cocaine and sucrose and saline control groups (F(_2,17_)=30.79, p<0.0001, followed by Tukey’s multiple comparison Test; sucrose vs. cocaine, p<0.0001; sucrose vs. saline, p=0.7874; cocaine vs. saline, p<0.0001; **Figure 3F**). Therefore, although sucrose-SA rats showed robust behavioral recall, only the cocaine recall group showed increased expression and altered localization of RyR2P.

As added controls, we also performed the following experiments. Using FIJI software, we reexamined the same images sampled above and increased the light intensity to saturation levels. However, we found that this only increased the background signal and signal in the dendrites. Additionally, using different slices of the same sucrose/cocaine recall animals (**Figure 2 B2-B3**), we examined the staining of pyramidal neuron cell bodies and dendrites in the motor cortex (M1) and hippocampal (CA1) regions (**Figure 3, supplemental figure 1A,B,** *left*). Relative to nondrug recall, we again observed a decrease in dendritic RyR2P staining with cocaine recall (**Figure 3, supplemental figure 1A,B,** *right*). Although the reductions in the M1 and CA1 regions were less robust than in the PFC, the general findings corroborated our previous results and indicated that cocaine recall likely impacts dendritic RyR2P in other regions of the brain as well.

### RyR2P during withdrawal or active drug-taking

Given the above results, we tested whether withdrawal caused changes in RyR2P during cocaine recall. We selected cocaine-SA and (control) sucrose-SA rats with 14 sessions of stable behavioral responding (**Figure 2 A,B2,B3**). Then, on day 15 at 24 hours after their last SA session, we perfused the animals, sectioned the brains, and stained for RyR2P and NeuN. Using the same antibodies and confocal settings as above (**Figure 3)**, we aligned the vmPFC sections by cell layer (**Figure 4A**) and compared RyR2P between the withdrawal groups. Photomicrographs showed robust expression of RyR2P in the apical dendrites (**Figure 4B**, *low magnification*) innervating L2-3 in all animals. Furthermore, a comparison of the somatic L5 staining between the groups (**Figure 4C**, *high magnification*) revealed no significant differences in RyR2P expression between cocaine and sucrose withdrawal groups (unpaired t-test; p=0.1691, t(6)=1.563, **Figure 4D**), indicating that withdrawal was not a driving factor in RyR2P expression.

**Figure 4.**
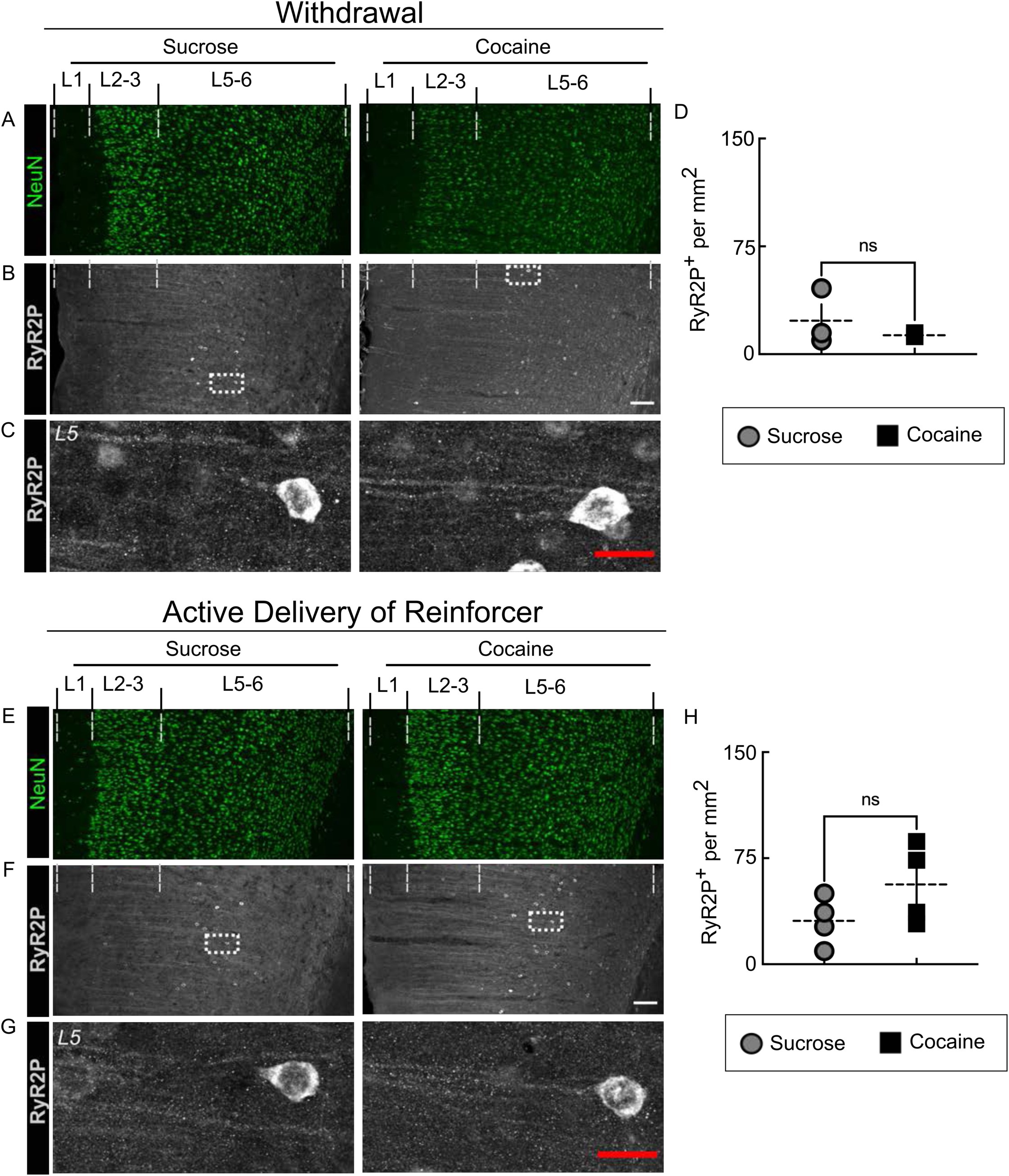
Spatial localization of RyR2P in vmPFC following withdrawal and self-administration. Experiments compared the expression of RyR2P in the vmPFC of sucrose-SA (*left*) and cocaine-SA (*right*) rats following either a 24h withdrawal (*top*) or active delivery (*bottom*) of the reinforcer. All animals were sacrificed on day 15 and the vmPFC brain sections were stained with antibodies against NeuN (**A,E***; green*) and RyR2P (**B,C,F,G**; *white*). The resulting confocal photomicrographs were collected at either low magnification (**A,B,E,F**; 10x) or high magnification (**C,G**; 60x) and aligned according to cell layer (L1-6). Summary bar graphs show the quantification of RyR2P positive neurons per mm^2^. No significant differences were detected comparing the sucrose and cocaine groups following withdrawal (**D**; unpaired t-test; p=0.1691, t(6)=1.563) or active delivery (**H**; unpaired t-test; p=0.4194, t(4)=0.8991). Scale bars: *white* = 100 μm; *red* = 20 μm.

As a follow-up, we also evaluated the opposite condition, where reinforcers were actively delivered. In a separate group of cocaine/sucrose-SA rats with 14 sessions of stable behavioral responding (**Figure 2 A,B2,B3**), we removed animals from an *ongoing* SA session (day 15) with light/tone cues after 90 minutes of reinforcer consumption and processed their tissue for comparison as above (**Figure 4E)**. In particular, photomicrographs from these two groups of animals also showed robust RyR2P expression in pyramidal cell apical dendrites (**Figure 4F,G**, *low and high magnification*), without significant differences in somatic L5 staining between groups (unpaired t-test; p=0.4194, t(4)=0.8991, **Figure 4H**, left). These data showed that alone, neither cocaine (withdrawal or activation) nor discrete cues (light, tone) were sufficient to alter RyR2P expression. Instead, the changes in RyR2P expression required a recall experience associated with cocaine.

### Co-labeling of RyR2P with Fos ensembles

PFC neurons with high Fos induction typically form ensembles of cells with highly correlated activity and higher sensitivity to environmental cues than non-Fos-induced cells (as reviewed in[30]). Using fluorescent immunohistochemistry, we determined whether vmPFC neurons with enhanced RyR2P expression had reliable Fos induction and whether this relationship differed between recall for drug and nondrug rewards. For this, we selected control recall rats with a history of yoked-saline or nondrug (sucrose-SA) (**Figure 2 B1,B2**) and a second group of drug recall rats (cocaine-SA or morphine-SA) (**Figure 2 B3,B4**). We reasoned that the inclusion of opioid morphine may provide an additional point of comparison for RyR2P. After double-label staining was complete, we imaged the vmPFC sections on a confocal microscope.

Photomicrographs (**Figure 5A**) of the control and drug recall groups showed somatodendritic expression of RyR2P surrounding Fos-positive nuclei (Fos+) and co-labeled RyR2P and Fos (RyR2P+Fos+) neurons. Therefore, for further comparison, we quantified the respective number of positive cells per mm^2^ section and analyzed the recall results. A one-way ANOVA indicated different levels of Fos expression (**Figure 5B**; F(_3,23_)=10.88, p=0.0001), and Fos+RyR2P+ (**Figure 5C**; F_(3,23)_=20.26, p<0.0001). A post hoc Tukey’s multiple comparison test found no differences in these three measures between the control yoked-saline and sucrose-SA recall rats (**Figure 5, supplemental figure 1, Table 1**). However, compared to control recall (yoked-saline or sucrose-SA), the drug recall (cocaine-SA or morphine-SA) (**Figure 5B,C,D**) showed increased expression of Fos+RyR2P+ (**Figure 5, supplemental figure 1, Table 1**). That is, relative to nondrug rewards, the recall of drugs significantly increased the percentage of RyR2+ labeled cells that co-labeled for Fos.

**Figure 5.**
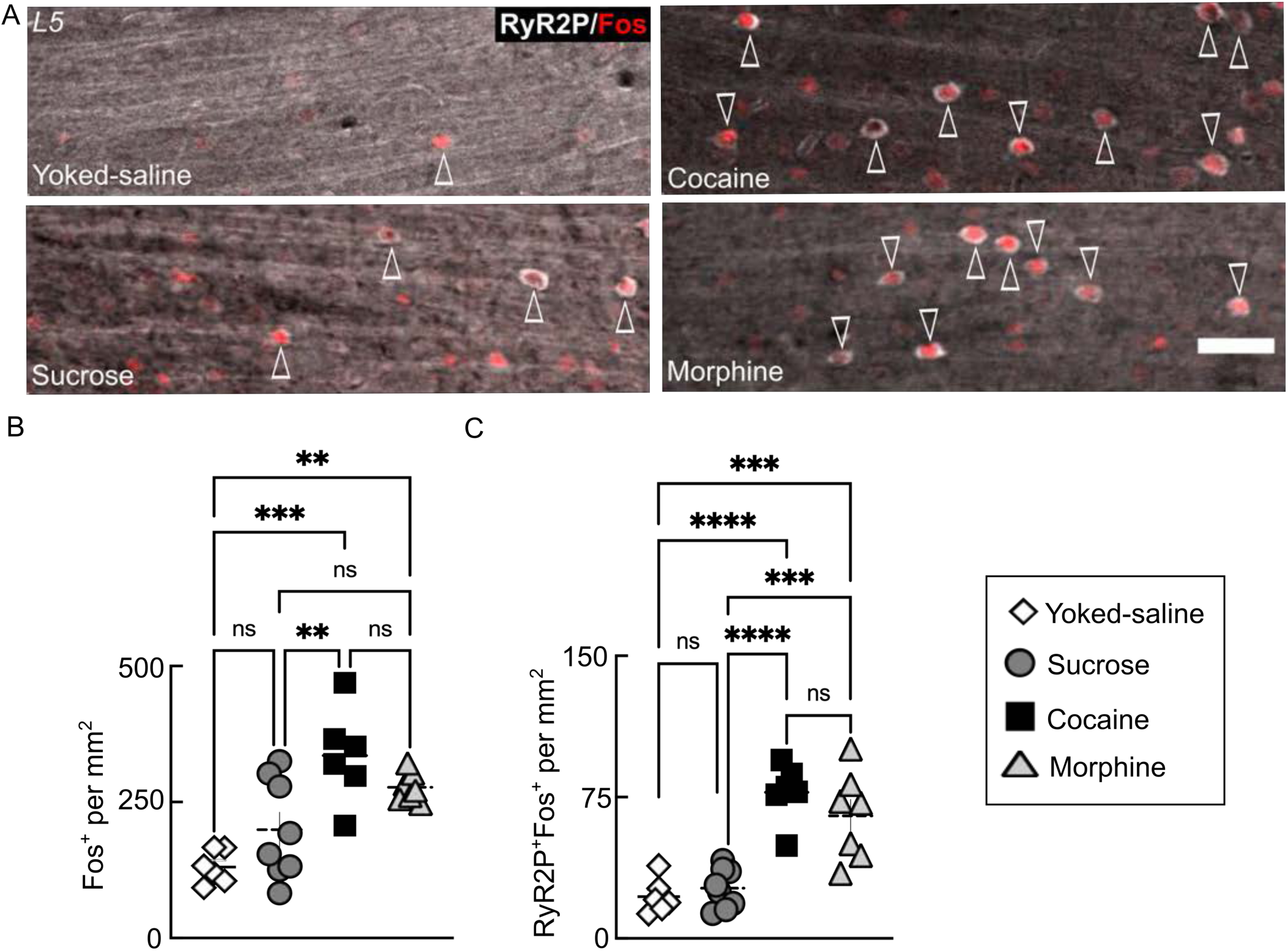
Increased colocalization of vmPFC RyR2P and Fos after cocaine or morphine recall. (**A**) Confocal photomicrographs of a stack of serial vmPFC sections showing differences in RyR2P (*white*) and Fos (*red*) colabeling (*white arrowheads*) in drug and nondrug recall tissue. (**B-D**) Graphs summarizing vmPFC differences in the expression of (**B**) Fos+ (F_(3,23)_=10.88, p=0.0001), and (**C**) RyR2P+Fos+ neurons (F_(3,23)_=20.26, p<0.0001). Note the enhanced numbers of RyR2P+Fos+ cells in drug reward (cocaine and morphine) relative to a nondrug reward (sucrose). Error bars represent S.E.M. One-way ANOVA and Tukey post hoc (*=p≤0.05, **=p≤0.01, ***= p≤0.001, ****=p≤0.0001). N=6-8 rats per experimental group with 4 technical replicates per rat. Scale bar = 50 μm

We conducted two additional control experiments measuring Fos, RyR2P+, and Fos+RyR2P+. The first experiment examined the dorsal medial PFC (dmPFC) region (**Figure 5, supplemental figure 2A**), where opioid induction is reported to be weaker[71,72]. As in those previous studies, we observed lower staining of Fos+, RyR2P+ and Fos+RyR2P+ in dmPFC morphine-SA recall tissue than in the dmPFC or vmPFC cocaine-SA recall tissue **(Figure 5**; **Figure 5, supplementary figure 1 Table 1; and Figure 5, supplementary figure 2 B,C,D),** and resembled results from yoked-saline and sucrose-SA recall rats. The second experiment used repeated injections of corticosterone or yohimbine to mimic chronic stress, which can enhance cocaine craving and RyR2 hyperphosphorylation[15,73]. However, in rats treated with chronic (12-18d) daily injections of cocaine or aformentioned drugs commonly used as pharmacological stressors (corticosterone or yohimbine) (**Figure 5, supplementary figure 3A)**, we did not observe significant changes in vmPFC relative to injections of saline (**Figure 5, supplementary figure 3B**). These RyR2P results indicated that morphine recall produced negligible changes in the dmPFC region and, at least under the conditions used here, neither chronic stress-like conditions nor chronic (experimenter-delivered) cocaine increased vmPFC RyR2P expression.

### RyR2 protein expression levels and phosphorylation

In addition to modulation by kinases, RyR2 dysregulation can involve increased protein expression, resulting in increased Ca2+ release from stores[74,75]. Using recall tissue from sucrose-SA and yoked-saline rats (**Figure 2)** as controls, we determined whether cocaine recall altered TRyR2 expression, phosphorylation, or both. Using immunofluorescence, we confirmed that in the vmPFC, the pattern of total TRyR2 somatodendritic staining in the sucrose-SA recall (**Figure 6A**, *left*) was similar to the cocaine-SA recall (**Figure 6A**, *right*) and was not noticeably different from naïve rats (**Figure 1A**).

**Figure 6:**
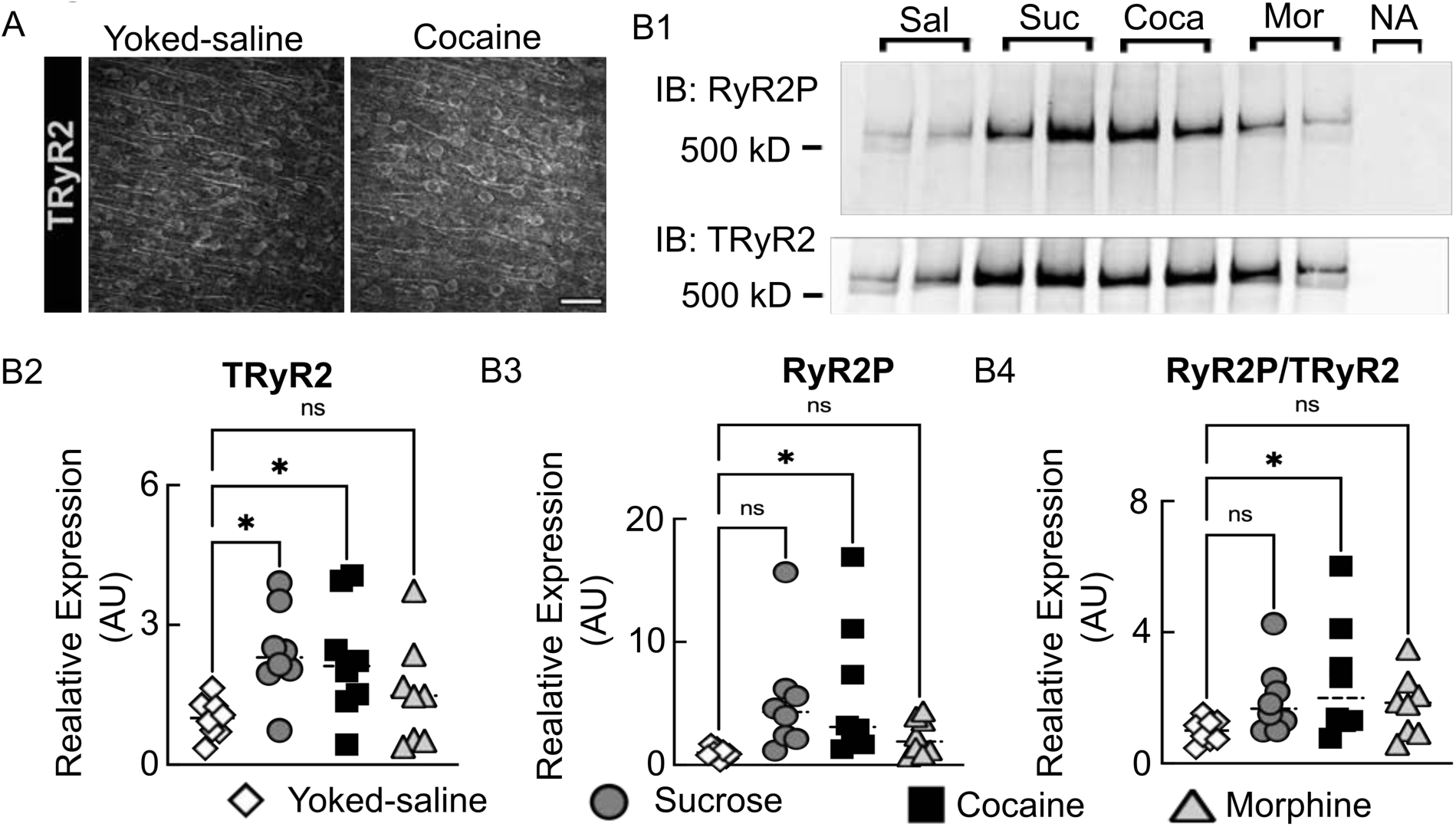
RyR2 expression and phosphorylation level following recall of natural and drug rewards. (**A**) 40x Confocal photomicrographs of vmPFC showing similar localization of TRyR2 (*white*) in yoked-saline and cocaine following recall. (**B1**) Representative Western blot of proteins pulled down using the anti-RyR antibody and probed using anti-RyR2P and anti-RyR2 antibodies. No antibody control (NA) shows no signal. (**B2**) Quantification of average TRyR2 (F_(3,28)_=3.497, p=0.0285), (B3) RyR2P (F_(3,28)_=3.171, p=0.0396) and (B4) total RyR2P normalized to the TRyR2 signal per experimental group. (F_(3,28)_=2.430, p=0.0861). One-way ANOVA followed by Dunnet’s multiple comparison post hoc *=p≤0.05. N=8-9 per experimental group, signal from 4 gels normalized to yoked-saline controls. Error bars represent S.E.M. Scale bar = 50 μm.

Next, using immunoprecipitation with a specific antibody against RyR (C3-33)[76], we immunoblotted with primary antibodies for RyR2P (**Figure 6B1,** *top*) and TRyR2 (**Figure 6B1,** *bottom*)[77]. We quantitated the expression of TRyR2, RyR2P, and RyR2P/TRyR2 (**Figure 6B2-B4)** and validated the efficacy of the immunoprecipitation by Western blotting. A one-way ANOVA indicated significant differences between animal groups (**Figure 6B2,** TRyR2: F_(3,28)_=3.497, p=0.0285; **Figure 6B3,** RyR2P: F_(3,28)_=3.171, p=0.0396; **Figure 6B4**, RyR2P/TRyR2: F_(3,28)_=2.430, p=0.0861). A post hoc Dunnett multiple comparison test revealed that the expression of total RyR2 (**Figure 6AB2**) in nondrug (sucrose, p=0.0221) and drug (cocaine, p=0.0459) groups was higher than in the yoked-saline group. However, RyR2P increased only with cocaine recall (**Figure 6B3** saline vs. sucrose, p=0.0753; saline vs. cocaine, p=0.0354) and RyR2P/RyR2 (**Figure 6B4** saline vs. sucrose, p=0.2518; saline vs. cocaine, p=0.0327). Although we quantified results from the morphine-SA recall tissue as well, sampling of the sensitive vmPFC region alone did not provide an adequate quantity of tissue to be useful for the pulldown assay. Therefore, the morphine results reflecting the combination of both sensitive (ventral) and insensitive (dorsal) PFC regions are difficult to interpret (TRyR2: morphine vs saline, p=0.5893; RyR2P: morphine vs saline, p=0.8163; RyR2P/TRyR2: morphine vs saline, p=0.4033; (also see **Figure 5, supplemental figure 2A;**[71,72]). Overall, these findings indicated that although recall of both sucrose-SA and cocaine-SA rewards increased TRyR2 protein levels, only recall of drug rewards were associated with hyperphosphorylation of RyR2.

### RyR2 transcription

RyR2 is also subject to activity-dependent transcriptional regulation[11]. To determine whether something similar occurred in vmPFC (**Figure 7 A**), we used *in situ* hybridization with RNAScope to label and compare RyR2 and Fos mRNA transcripts after drug and nondrug recall. Immunofluorescent photomicrographs of sucrose-SA recall (**Figure 7 B,** *top*) and cocaine-SA recall (**Figure 7 B,** *bottom*) showed overlapping expression of Fos and RyR2 mRNA with a high degree of colocalization within the nuclei (**Figure 7 B,** *right top and bottom*).

**Figure 7.**
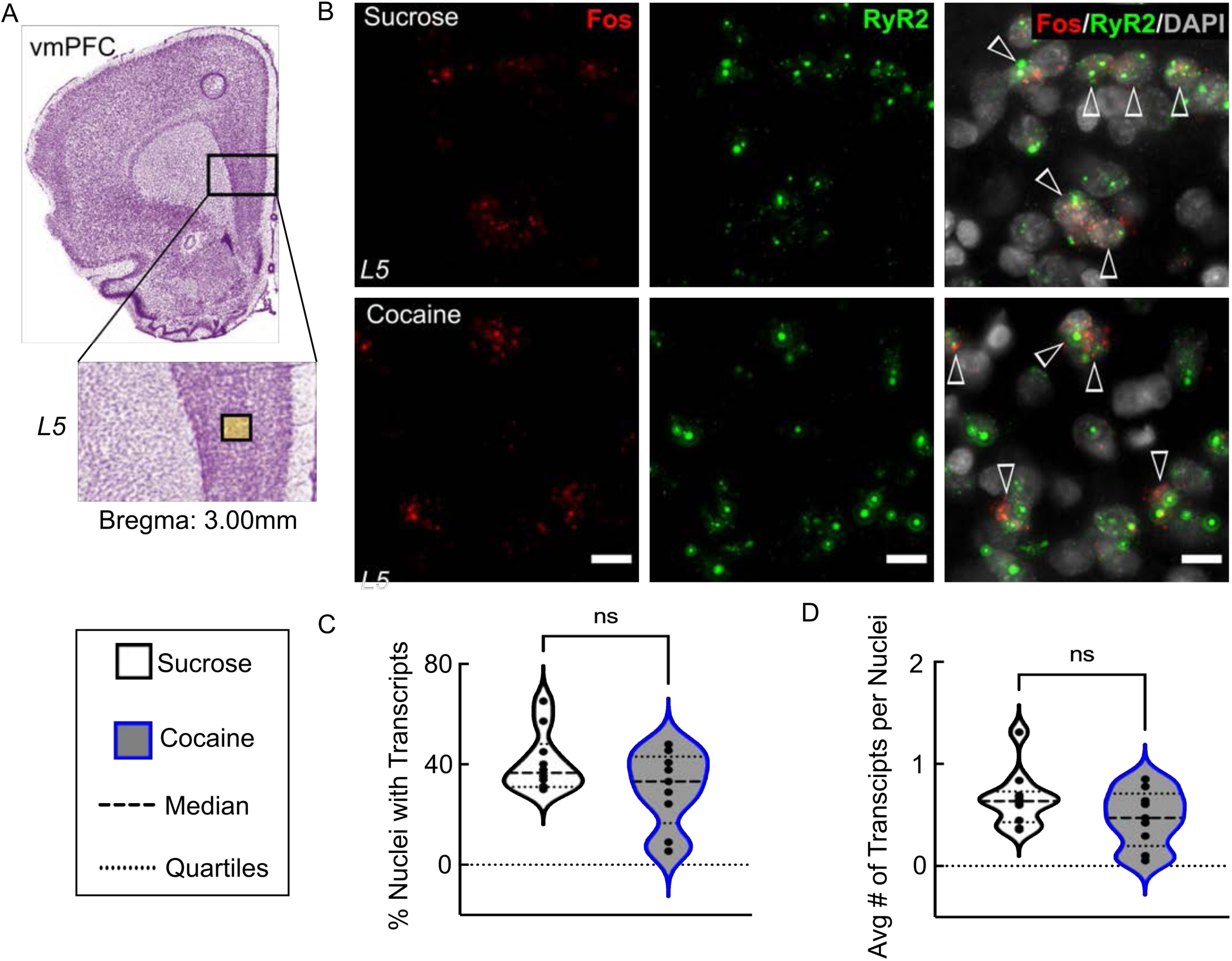
Recall of drug and natural rewards is associated with similar expression levels of mRNA for Fos and RyR2. (**A**) Representative confocal photomicrographs showing the L5 vmPFC site of analysis of (**B**) mRNA expression of Fos (*red*), RyR2 (*green*), DAPI (*grey*) or their co-expression (*white arrowheads*) in sucrose recall (*top*) and cocaine recall (*bottom*) tissue. (**C, D**) Summary quantification of RyR2 transcripts expressed as (**C**) the percent of nuclei with transcripts, (**D**) and the average number of transcripts per nuclei. A comparison of transcripts in sucrose and cocaine recall showed no significant differences. ns=p>0.05. N=3 rats per experimental group with 2-4 technical replicates per rat. Scale bar = 10 μm.

However, quantification and analysis of the RyR2 transcripts using an unpaired t-test revealed no significant differences between the recall groups. Results were comparable whether expressed as a percent of nuclei (**Figure 7C**; p=0.1147, t_(17)_=1.663) as the average number of transcripts per nuclei (**Figure 7D**; p=0.1585, t_(17)_=1.475). For technical reasons, we could only effectively quantitate the average number of RyR2 transcripts in nuclei coexpressing with Fos mRNA in tissue from two subjects. That analysis showed no statistical difference between sucrose and cocaine-SA (RyR2 transcripts in Fos+ p=0.2770, t_(13)_=1.135, mean of sucrose=3.312, mean of cocaine=2.701). Therefore, although drug recall increased protein levels of RyR2P relative to the total RyR2, we concluded that *RyR2* mRNA expression was unaffected, indicating that drug recall did not alter RyR2 transcription.

## Discussion

RyR2 is expressed in the heart and multiple brain regions[9,78]. Recent work associates pathological changes at the S2808 site of RyR2 function with excessive phosphorylation and cognitive deficits[15,79,80]. Although the expression of hippocampal RyR2 has been extensively studied, its role in the PFC and cognitive dysfunction is largely unknown. In the present study (summarized in **Figure 8**), we determined the subcellular distribution of PFC RyR2 and then evaluated the impact of behavioral conditioning (reward-SA) on its expression and phosphorylation. Under basal conditions, vmPFC soma and dendrites expressed tRyR2, but we only detected phosphorylated S2808-RyR2 in the apical dendrites of pyramidal cells. Following operant conditioning, we also found that recalling the memory of a previous reward (drug or nondrug) increased the overall expression of tRyR2 protein. However, when the memories were associated with addictive drugs, the phosphorylated S2808-RyR2 (RyR2P) increased and RyR2P underwent physical reorganization from dendrites to activated Fos+ neuron somata. These findings provide new insights into the role of RyR2 phosphorylation in decision-making for ordinary rewards as well as a phosphorylation-based post-translational modification relevant to drug seeking.

**Figure 8:**
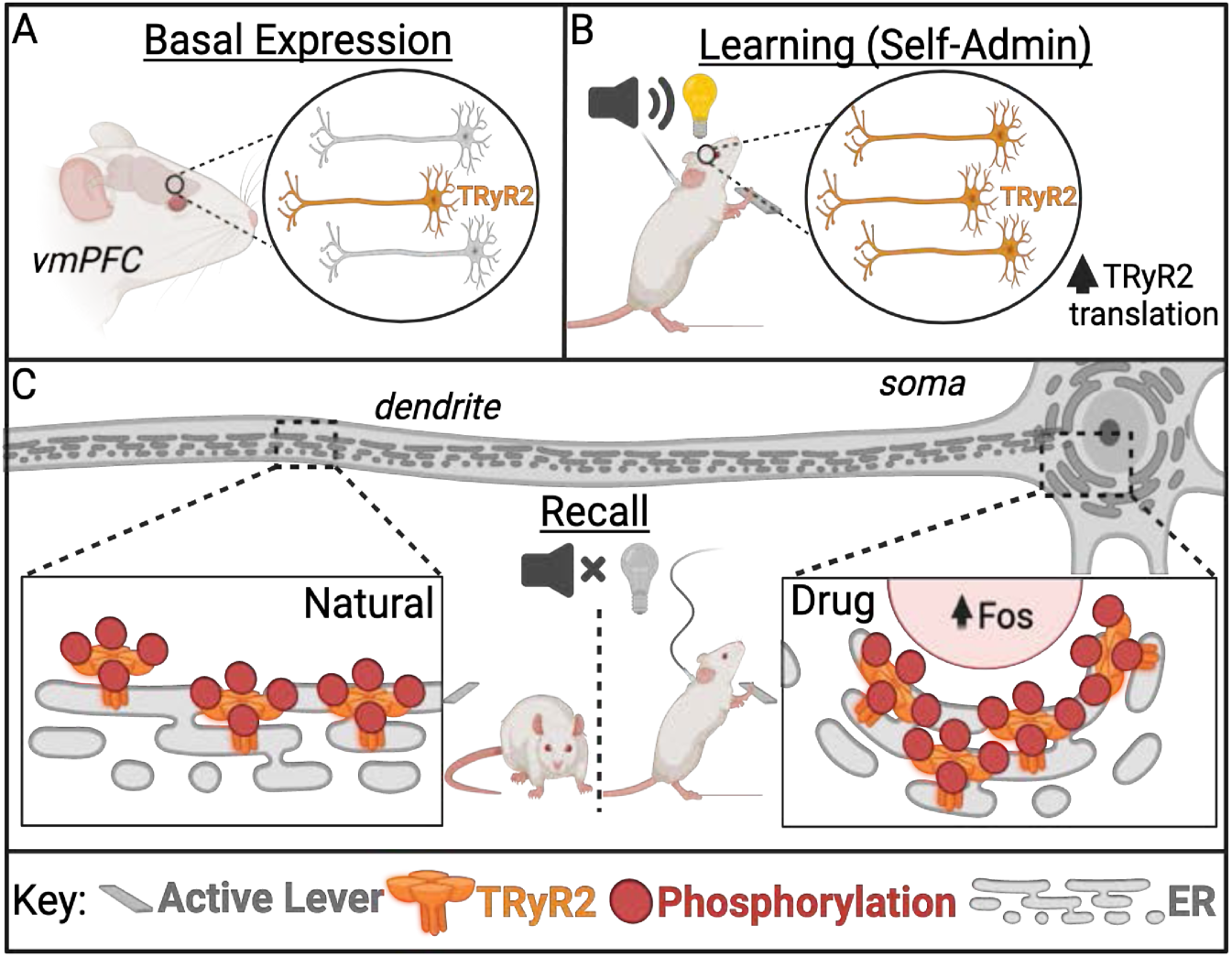
Key Findings. A) Under basal conditions, TRyR2 was expressed in the somata and apical dendrites of L2-3,5 pyramidal cells in the vmPFC. (B) The expression of TRyR2 protein increased after SA training for nondrug (sucrose) and drug (cocaine or morphine) rewards. (C) The expression of phosphorylated S2808-RyR2 (RyR2P) in nondrug (sucrose, *left*) and drug (cocaine or morphine, *right*) rewards differed. RyR2P expression in the apical dendrites in nondrug recall resembled that observed in naïve rats (*left inset*). In contrast, in drug recall (*right inset*), RyR2P was no longer detected in the apical dendrites and was instead expressed in soma colocalized with Fos, a marker of highly activated neurons. Made with BioRender.

### Basal arrangement of TRyR2 & RyR2P

In addition to the hippocampus[9,78], RyR2 is also expressed in the cerebral cortex[2,81,82]. However, less is known about the subcellular distribution of the RyR2 isoform in mPFC and its function in these neurons classically associated with decision-making, memory, and task engagement[83]. In addition to its role in cognitive control processes, vmPFC function is negatively impacted by diseases such as addiction[84]. Although anti-RyR2 antibodies have been used to survey RyR2 in brain and peripheral tissue[2,81,82], the broad scope of the studies and the unclear characterization of the antibodies made it difficult to infer their impact on related behaviors. Therefore, we performed a colocalization analysis with known neuronal subcellular markers in fixed brain slices, and ultimately identified RyR2 in the soma and apical dendrites of CaMKII-positive pyramidal neurons in cell layers 2-5. Although a previous striatal study suggested that RyRs regulate excitability in GABAergic neurons in other regions[85], we did not regularly observe TRyR2 expression in GABAergic PV/SST interneurons.

As phosphorylation of S2808 is a common post-translational modification of RyR2, we characterized a phospho-specific 2808 antibody and directly visualized the vmPFC. We found RyR2P in apical dendrites of L5 pyramidal neurons, but not typically in vmPFC soma. The unequal distribution of RyR2P was unexpected and significant, as apical dendrites represent a distinct distal compartment of the dendritic tree and serve as the postsynaptic site for most excitatory glutamatergic synapses[86]. Although this expression pattern has not been previously reported in neurons, basal phosphorylation of S2808-RyR in cardiac cells occurs in multiple animal species and humans[17–20]. Therefore, phosphorylation of RyR2 by CaMKII or PKA may significantly alter channel function in pyramidal neurons like that observed in the heart, including increased open probability (Po)[21,87], enhanced channel activity (Po) in response to steep changes in Ca^2+^[88], and dissociation of regulatory factors (eg, FKBP12.6)[21].

### Reward recall increases TRyR2

Success in spatial or fear memory tasks requires associative learning and increases in TRyR2 expression[11,89,90]. How associative learning impacts PFC RyR2 is largely unexamined. To this end, we employed a recall test that engages vmPFC[69,91] following an instrumental learning task (daily SA-training), where rats learned to differentiate between choice behaviors (active vs. inactive lever) for a reward. Then, after recall testing, we assayed TRyR2 as in previous studies [15,92], using pulldown Western blots and direct single-molecule visualization of individual neurons by RNAscope. Our results showed that independent of the type of reward, recall increased TRyR2 protein content without obvious differences between rewards in RyR2 mRNA. Although recall testing occurred the day after SA, we acknowledge that the half-life of RyR2 protein is long (∼8d [93]). We cannot rule out the possibility that the increased TRyR2 reflected the original SA training. Nor have we examined the additional potential influence of degradation mechanisms regulating TRyR2 protein or mRNA[94–98]. In addition, other studies (below) indicate the increases in TRyR2 protein may be task-specific.

Several previous studies investigated hippocampal RyR2 using a Morris water maze as a measure of spatial memory important for remembering and navigating through space or contextual fear conditioning as a measure of emotional memory that employs conditioning to associate a context and a cue with a brief stimulus. Whereas spatial memory tasks increased RyR2 mRNA and protein for more than 10 days[11,89], fear memory produces transient increases in RyR2 protein expression (<29 hr)[90]. Subsequent brief re-exposure (that is, minutes) to the same fear context (without reinforcement) acutely increased TRyR2 protein levels as well[90]. Although these were hippocampal studies, the design of our PFC recall task (brief context re-exposure without reinforcement) resembled the fear memory extinction training study, bolstering our supposition that recall increased TRyR2 protein translation. The results of these present and past studies indicate that rather than the context, increases in RyR2 require associative learning[90,99] and that RyR2 mechanisms are highly responsive and adaptable to a specific task.

Other existing studies show that single or repeated injections of nicotine[13] or methamphetamine[41] also increase RyR2 mRNA or protein in brain regions, including the frontal cortex. Although this work highlights the general sensitivity of RyR2, the specific role of RyR2 in drug self-administration and the recall/seeking process is unclear. Just as drug-SA and recall activate different vmPFC ensembles[65], SA and noncontingent (experimenter-delivered) drugs can activate different brain circuitry [100–105]. So, it is uncertain whether the previously published effects of nicotine and methamphetamine reflect the drug *per se* or a conditioning process. Alternatively, the intensity of acute high-dose drug stimulation may play a role, as brief theta-burst stimulation[89] increases TRyR2 protein expression as well. Another caveat of all biochemical approaches, including our own, is that microglia and blood vessels also express RyR2[106,107]. Although our IHC neuronal staining corroborated our cocaine Western blot findings, we did not measure neuronal Ca^2+^ within neurons. Perhaps our strongest support for RyR2 neuronal activation was the use of low-dose daily SA and the delayed recall test, which showed differences between reward and yoked-saline rats that lacked a reinforcer but had similar behavioral experiences.

### Drug recall expands RyR2P/Fos(+) ensembles

Although the increase in TRyR2 expression was comparable in nondrug and drug recall, the degree of phosphorylation (RyR2P) was not. The localization of RyR2P with Fos+ neuronal ensembles has important implications for the role of RyR2 in substance use disorders (SUD). Although previous studies indicated that reexposure to reward environments increases Fos expression in PFC[69,108–113] and certain Fos+ recall ensembles are necessary for drug-seeking[69,91], how and why vmPFC circuitry is reorganized by addictive drugs remains unclear. The elevated RyR2P+Fos+ response in the cocaine and morphine rats actively seeking drug implies a direct role for RyR2P in compulsive behaviors. On the contrary, the low levels of RyR2P+Fos+ neurons with sucrose recall were indistinguishable from yoked-saline rats, indicating that increased RyR2P was not required for memories related to ordinary natural rewards. However, more than simply a drug response, RyR2P activation likely depends on the learned association, as we did not observe increased RyR2P with stress-mimicking drugs or cocaine withdrawal or cocaine delivery under non-contingent conditions which likely did not evoke similar memories. Instead, the increase was apparent with drug recall only when the expected associated reward response was being actively pursued.

In line with this, Fos studies show that drug and nondrug recall activate different Fos+ ensembles[69,114] just as positive and negative experiences are represented by different PFC populations[115,116]. Therefore, Fos+ ensembles previously reported to drive cocaine/opioid seeking are likely attributable to increased phosphorylation at S2808-RyR2 (RyR2P) rather than simply increased TRyR2 expression. To the best of our knowledge, this has not previously been reported. It is important to note that, for the sucrose recall experiments, we could not internally corroborate our all of our IHC findings with Western blot. Alone, the ventral PFC provided insufficient tissue for the analysis of morphine recall and similar to previous opioid reports[117], we confirmed that Fos/RyR2P in dorsal PFC was unchanged by morphine recall. Differences between ventral and dorsal PFC responses are discussed in detail elsewhere[114,118].

### Drug recall stimulates RyR2P reorganization

Alterations in phosphorylation can influence RyR2 channel distribution and its coordinated level of activity[119]. How phosphorylation impacts RyR2 in PFC is largely unexplored. We found that low level phosphorylation of S2808-RyR2 (sucrose-recall) supported dendritic localization, consistent with myocyte studies showing low level PKA stimulation increases RyR2 clusters[120] and enhances the propagation of Ca2+ waves[121–123]. But drug recall was associated hyperphosphorylation that significantly impaired the usual distribution of RyR2P in apical dendrites. Thus, the underlying changes likely involve the drug-related hyperphosphorylation, which resembles cardiomyocyte studies showing prolonged PKA stimulation has an opposite effect of dispersing RyR2 clusters, slowing Ca2+ waves and increasing ‘silent’ leak Ca2+ in remaining RyR2 channels[119,124]. Reductions in RyR2 activity also diminish the inhibitory slow afterhyperpolarization current (IsAHP) in hippocampal dendrites[125,126], resulting in compensatory neuronal excitability[127,128]. We previously reported reduced IsAHP currents in PFC Fos+ neurons from rats recalling and actively seeking cocaine as well[62].

Alongside the dendritic reduction, we found that cocaine and morphine recall stimulated RyR2P expression in Fos+ soma. Cocaine and morphine share the common end result of superactivating adenylate cyclase/cAMP/PKA signaling pathways associated with addiction[129–133]. Given the critical role of PKA in learning and neuronal excitation, our data suggested that enhanced somatic RyR2P in Fos+ neurons may result in increased drug seeking by reducing the IsAHP and facilitating the learned association with the drug. Interestingly, in hippocampus, RyRs couple to and are activated by L-type Ca^2+^ channels (LTCCs)[134,135] clustered on the plasma membrane. Together, these two Ca2+ sources activate K+ channels that regulate the IsAHP[125] and contribute to transcription factor activation including CREB-dependent gene expression[136,137]. However, precisely how increased somatic RyR2P impairs the IsAHP remains to be determined. Our previous work with cocaine-SA, extinction and reinstatement showed desensitization of IsAHP, but we did not test the direct involvement of RyR2P-Ca2+ and inhibitory calmodulin for instance[62]. Interestingly, RyR2 amplification of activity-dependent calcium influx stimulates hippocampal CREB phosphorylation and enhances the expression of transcription factor Npas4[12,138]. Furthermore, CREB plays a role in the induction of immediate-early genes including *c-fos* (reviewed in [139]). Thus, similar alterations may also influence membrane excitability and gene expression in PFC pyramidal cells after chronic drug-SA and recall.

In summary, in the present study, we demonstrated that TRyR2 is localized throughout the soma and dendrites of pyramidal cells and, under basal conditions, constitutively phosphorylated at S2808 in apical dendrites. We also showed that drug and nondrug reward recall generally increase TRyR2 content. However, drug recall was unique in potentiating RyR2P, impairing the usual dendritic expression and robustly stimulating somatic localization in Fos+ neurons, classically associated with increased neuronal excitability and impaired choice behaviors. This work provides new insights into RyR2 neuroadaptations associated with drug-seeking behavior.

## Methods

### Self-administration

#### Subjects

Male Sprague-Dawley rats 250-275g (n=73) sourced from Envigo Corporation were individually housed on a 12-hour reverse light/dark cycle with Ad libitum access to water. Daily food intake (restricted to 10-20g) occurred after operant sessions. A subset of this group was used for immunofluorescent and RNAscope studies (total n = 41) studies or immunoprecipitation and immunoblotting experiments (n = 32).

#### Intravenous surgeries

Using our published procedures[62], we anesthetized rats with isoflurane gas (induction 5%; maintenance 2.1-2.5%) and inserted a silastic catheter (22-gauge) into the right jugular vein prior to externalization and attachment to a vascular access button (Instech, VABR1B/22). Rats received ketorolac (2 mg/kg, IP) following surgery. During homecage recovery (4-7d), catheters were flushed daily with heparinized saline containing 8mg/kg of gentamicin (Henry Schein, 1098195). After daily SA training, catheters were flushed with 0.1ml of a sterile saline solution containing 500 units/ml of heparin, followed by a 0.1ml heparin lock solution (300 units/ml).

### Behavioral procedures

#### Self-administration training of sucrose, cocaine, and morphine

As described previously[62,63], we trained rats on a fixed-ratio-1 (FR-1) schedule in a 2-hour (cocaine, yoked-saline) or 3-hour (morphine, sucrose) session. Active lever presses resulted in the release of a sucrose pellet (45mg), 100µl infusion of cocaine (0.5mg/kg per infusion), or 100µl infusion of morphine (0.5mg/kg per infusion). Rat dosing was calculated from daily weight measurements, and drug (cocaine HCL (NIDA, 2.5 mg/kg) and morphine sulfate pentahydrate (NIDA, 2.5 mg/kg) were prepared weekly in sterile saline. Summarized responses (total number of infusions and lever press activity) are shown in **Figure 2**. Three rats were excluded from SA due to surgical complications or failure to learn the behavior.

#### Recall, withdrawal, and active drug-taking

To create an extinction-like condition associated with reward-seeking, recall tests (1.5h) were conducted in the SA boxes without rewards or discrete cues as described previously[30,109]. As an additional control, a cohort of cocaine-SA and (control) sucrose-SA rats received either withdrawal (no reward or discrete cues) or active drug-taking test (with reward and discrete cues) on their last session day (**Figure 2A**). Rats were sacrificed immediately after testing by rapid decapitation or cardiac perfusion with PBS and 4% PFA. An unpaired t-test was used to determine the difference in inactive and active lever presses during the recall test.

#### Non-contingent cocaine and stress treatments

Male Sprague-Dawley rats (Envigo Corp.) 300-350g (n=12) were randomly assigned to one of five treatment groups to receive chronic (12-18d) daily treatments of: saline/saline (n = 3), vehicle/cocaine (n = 3), yohimbine/saline(n = 3), corticosterone/saline (n = 3), or yohimbine/saline (n = 3). Dosing was performed by intraperitoneal (IP) injections (separated by 15 min) delivered to opposite sides at a volume of 1ml/kg at the following doses: corticosterone (2 mg/kg), cocaine (20 mg/kg), yohimbine (2 mg/kg), or vehicle (equal volume of 10% EtOH in distilled water). Stock solutions of cocaine HCl (NIDA, 20 mg/kg) were prepared weekly in sterile saline, while corticosterone (Sigma Life Sciences 27840, 2 mg/kg) and yohimbine hydrochloride (Sigma Life Sciences Y3125, 2 mg/kg) were prepared daily in a 10% ethanol (EtOH) solution in water.

### Molecular Endpoints

#### Constructs and Transfection

Previous studies evaluated the specificity of a phospho-antibody directed against RyR2P **(Figure 1-figure supplement 2**) [51,52]. As an additional control, we transfected human embryonic kidney cells with wild-type RyR2 (WTS2808) plasmids or a mutant RyR2 S2808 converted to an alanine (MTS-A2808) that prevents phosphorylation. We used double-label fluorescence immunohistochemistry to evaluate RyR or colabeled RyR2P-immunoreactive cells, with or without forskolin (10 µM; 1 hr), to activate adenyl cyclase and PKA [53].

#### Immunofluorescent Staining

At 90 min after testing, rats from the recall (n=27), drug on board (n=8) and withdrawal (n=6) groups were anesthetized with isoflurane and perfused with 100 ml of 0.1 M phosphate-buffered saline (PBS), followed by 400 ml of 4% paraformaldehyde in PB solution. Brains were post-fixed in paraformaldehyde (24 h) and transferred to a sucrose (30%) PBS solution at 4°C for 2 days. Coronal brain sections (40 um) of the vmPFC (A32V)(bregma: +3.00 to +3.72 mm; Paxinos & Watson V7, 2013) were obtained using a microtome and selected for staining.

The free-floating sections were washed and blocked with a solution containing 3% normal donkey serum (NDS) and 3% bovine serum albumin (BSA) in PBS, supplemented with 0.3% Triton X-100 and 0.2% Tween 20, for a duration of 2 hours. Following blocking, sections were incubated overnight at 4°C with primary antibodies **(Table 3**, below) diluted in the same blocking solution. Sections were incubated for 1 hour with secondary antibodies, diluted in PBS containing 3% NDS and 3% BSA, supplemented with 0.3% Triton X-100 and 0.2% Tween 20.

**Table 3:**
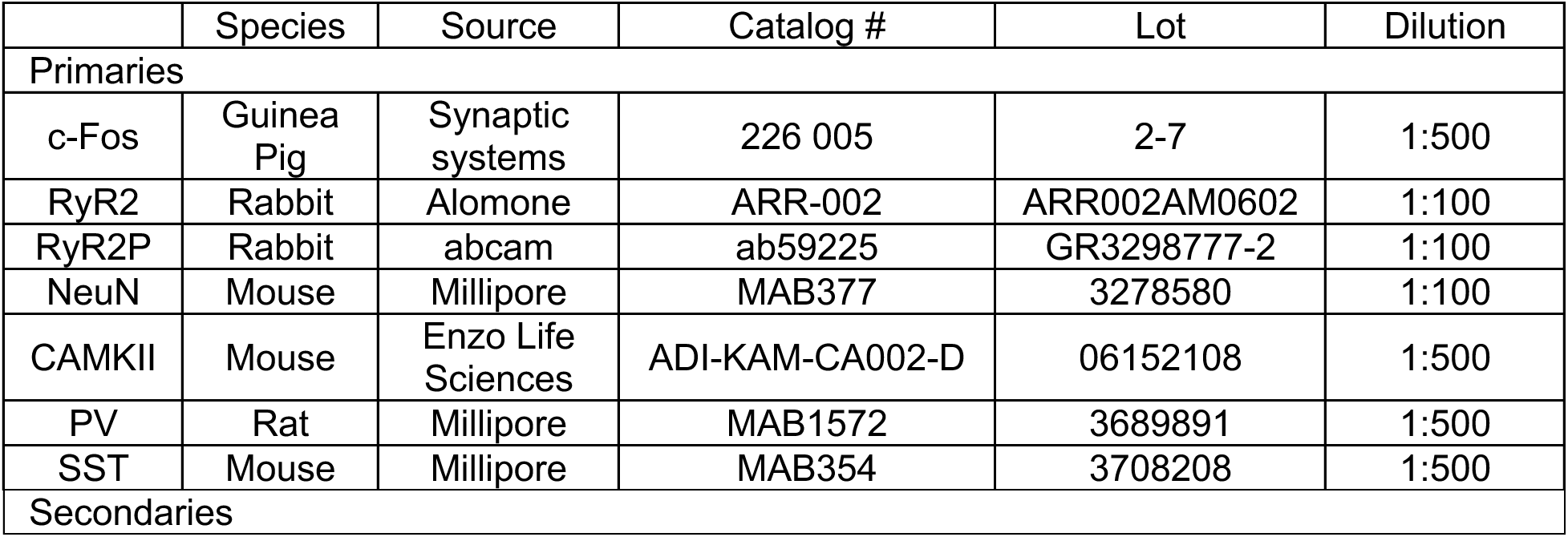
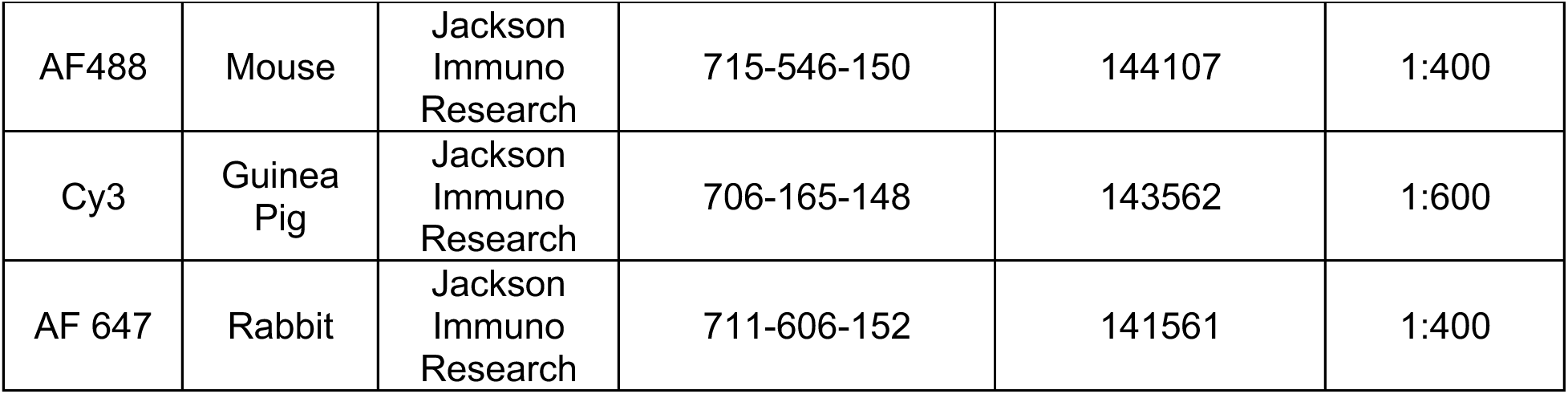
List of primary and secondary antibodies used for immunohistochemistry.

Following DAPI staining, the sections were mounted on Fisher Superfrost Gold Plus slides (cat #22-035813) and coverslipped using Prolong Glass Antifade media (Invitrogen, P36980).

#### Imaging and Analysis

Imaging of the sections was conducted using an Olympus Fluoview FV1200 microscope, with images captured at magnifications of 10×, 20×, 40×, or 60×. Consistent equipment and software settings were maintained throughout the duration of the experiments. The areas of interest within the vmPFC (A32V) were selected based on a rat brain atlas (Paxinos & Watson V7, 2013), and the corresponding locations were indicated by overlays on schematic brain coronal sections (**Figure 3**). The quantified areas encompassed layers 2 to 6 of the prefrontal cortex and measured 0.72 mm². Fos+ nuclei were quantified on a single focal plane within the z-series using the Find Maxima software in FIJI. Fos+RyR2P+ per mm² values were calculated manually using the FIJI multipoint selection tool. For each rat, the values of Fos+ and Fos+RyR2P+ per mm² were obtained from two sections (4 hemisphere images) and averaged, resulting in a single n value per brain area. Statistical analysis was conducted using one-way ANOVA followed by Tukey’s post hoc test to determine differences between groups.

TRyR2/CAMKII and TRyR2/PV/SST colocalization quantification used the JACoP plugin to calculate the Manders’ correlation coefficient. The thresholded Mander’s M value corresponds to the fraction TRyR2 overlapping with the neuron cell type marker (CAMKII, PV, or SST). One-way ANOVA followed by Tukey’s multiple comparison test was used to identify differences in colocalization.

#### Immunoprecipitation and Immunoblotting

Bilateral punches from the rat prefrontal cortex were lysed using a Polytron homogenizer in a lysis buffer containing 20 mM NaF, 10% glycerol, 50 mM Tris-Cl (pH 7.5), 150 mM NaCl, 0.5% NP-40, 1 mM PMSF, 1 mM EDTA, and 1X protease inhibitor cocktail (Fisher Scientific, #53-514-21SET). The lysates were solubilized on a rotator overnight at 4°C, and the supernatant was obtained by centrifugation at 20,000g for 10 min at 4°C. Immunoprecipitation of RyR2 was performed using 100-400 μg of protein per precipitation in a 200-300 μL volume. The lysates were incubated overnight at 4°C with a 1:100 dilution of anti-RyR2 antibody (C3-33, Invitrogen). Following the overnight incubation, Protein A-Sepharose beads (Fisher Scientific #PI20423) were washed with lysis buffer and added to the tubes at a volume of 30 μL per precipitation. The samples were incubated for 3 hours at room temperature on a shaker. Subsequently, the beads were washed four times with lysis buffer and spun at low speed (500g for 3 minutes) to collect the beads between each step. Pellets were cleared of buffer using a 30g needle. The samples were then heated in 45 μL of 1x LDS sample buffer at 95°C for 5 minutes, followed by centrifugation, and 40 μL of the lysate was removed. SDS-PAGE was performed using a 3-8% Tris Acetate gradient gel, and the separated proteins were transferred to a nitrocellulose membrane at 30V and 4°C for 1-3 hours. The membrane was then blocked with 5% nonfat dry milk in tris-buffered saline (TBS).

For immunoblotting, primary antibodies for RyR2P and TRyR2 (**Table 4**) were incubated overnight at 4°C in TBS containing 0.1% Tween 20 (TBST) and 5% fraction V bovine serum albumin. After three sequential washes with TBST, blots were incubated with secondary antibodies in 5% nonfat dry milk in TBST for 1 hour at room temperature. The images were captured using an Azure Sapphire imager, and the blot densities were analyzed using Azure Spot analysis software. One-way ANOVA followed by Dunnett’s multiple comparison test was used to identify differences between groups.

**Table 4:** List of primary, and secondary antibodies, and reagents used for immunoprecipitation and immunoblotting.

| Antibody | Species | Source | Catalog# | Lot# | Dilution |
| --- | --- | --- | --- | --- | --- |
| Ryanodine Receptor-2 (C3-33) | mouse | Invitrogen | MA3-916 | UK293889 | 1:100 |
| P-RYR-2 (phospho S2808) | rabbit | Invitrogen | PA5-36758 | WB3186373 | 1:500 |
| RyR2 | Guinea pig | Frontier Institute | FI-RyR2-GP-AF480 | MSFR105290 | 1:500 |
| HMW Ladder | NA | Life Technologies | LC5699 | 2441668 | NA |
| IRDye800 | Rabbit | LiCor | 926-32211 | -- | 1:5000 |
| IRDye800 | Guinea Pig | LiCor | 926-32411 | -- | 1:5000 |

#### RNAscope assay

RNAscope in situ hybridization (ISH) was performed on 2 rats per experimental group in coronal sections (40 um) from the vmPFC (Bregma: +3.00 to +3.72 mm) (Paxinos & Watson V7, 2013). Manual procedures were performed according to the User Manual for Fixed Frozen Tissue, utilizing the RNAscope Multiplex Fluorescence Reagent Kit V2 from Advanced Cell Diagnostics. For fluorescent labeling, sections were incubated with specific combinations of colors associated with each channel, such as orange (Opal 620) and far-red (Opal 690). Images were captured digitally at 40x magnification using a Zeiss LSM 880 microscope. RNAscope signal was quantified using dotdotdot software[140], an automated program that calculated the percentage of Fos+, RyR2+, and nuclei co-labeled with RyR2 and Fos. Additionally, the average number of RyR2 transcripts in Fos+ nuclei was quantified. Two sections (2-4 hemisphere images) per rat were counted. Statistical analysis was performed using an unpaired t-test to determine differences between groups.

## Acknowledgments

The authors thank Dr. Laurent Martin for RNAscope reagents and guidance, the Comprehensive Pain and Addiction Center (CPA-C), and the Center of Excellence in Addiction Studies (CEAS) NIH/NIDA DA051255 for their support. We would also like to thank Patty Jansma and the UA microscopy core for training and guidance with the LSM 880. The authors also thank Dr. Wayne Chen for kindly providing point constructs for our validation experiments.

---

**Figure 1, supplemental figure 1.**
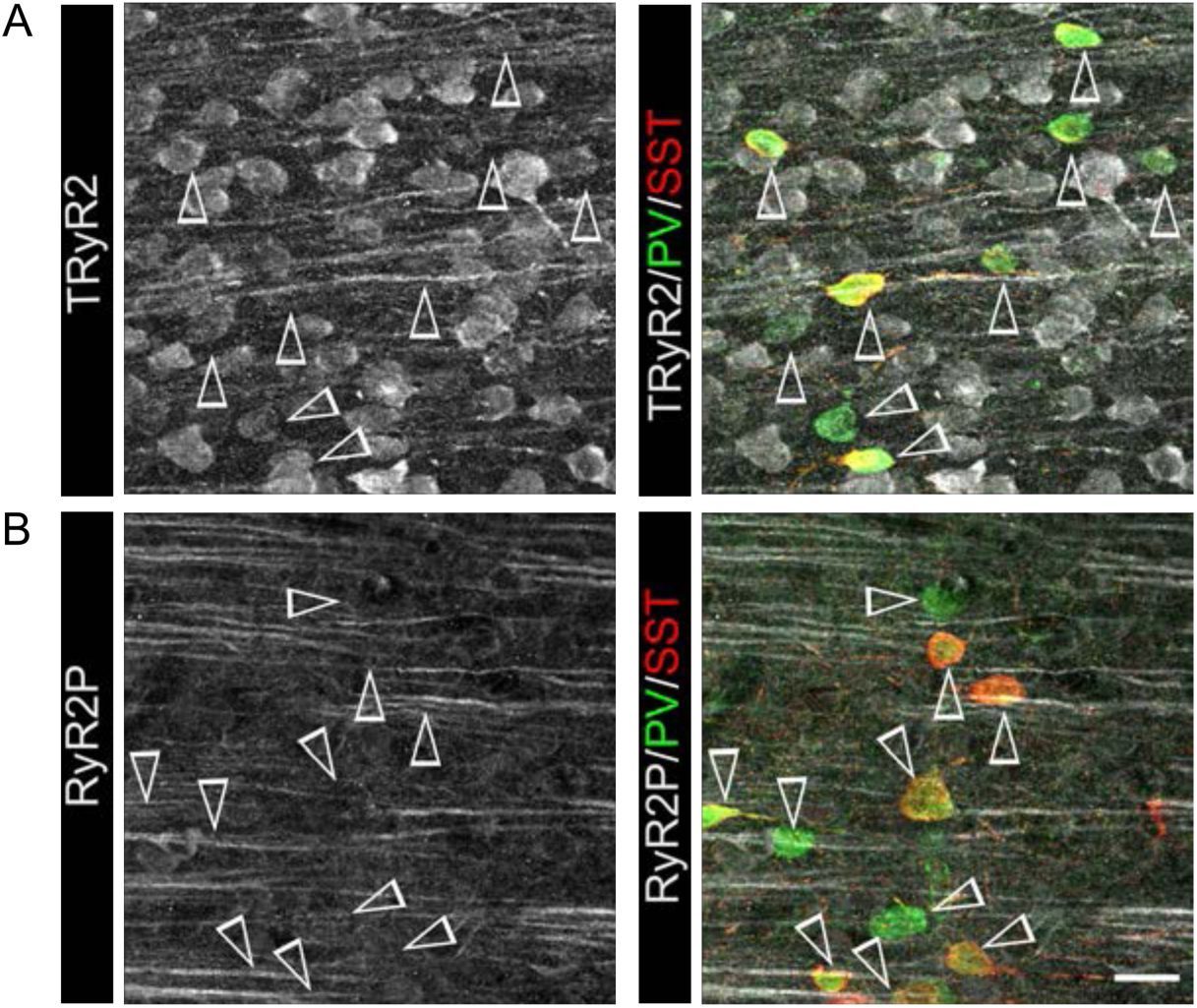
TRyR2 and RyR2P are not highly expressed in inhibitory neurons of the PFC. Representative confocal photomicrographs showing the low degree of colocation of (**A**) TRyR2 (*white*) and (**B**) RyR2P (*white*) with PV (*green*) and SST (*red*), two markers of inhibitory neurons in the PFC. Note: White open arrowheads indicate the location of PV and SST positive neurons.

**Figure 1, supplemental figure 2.**
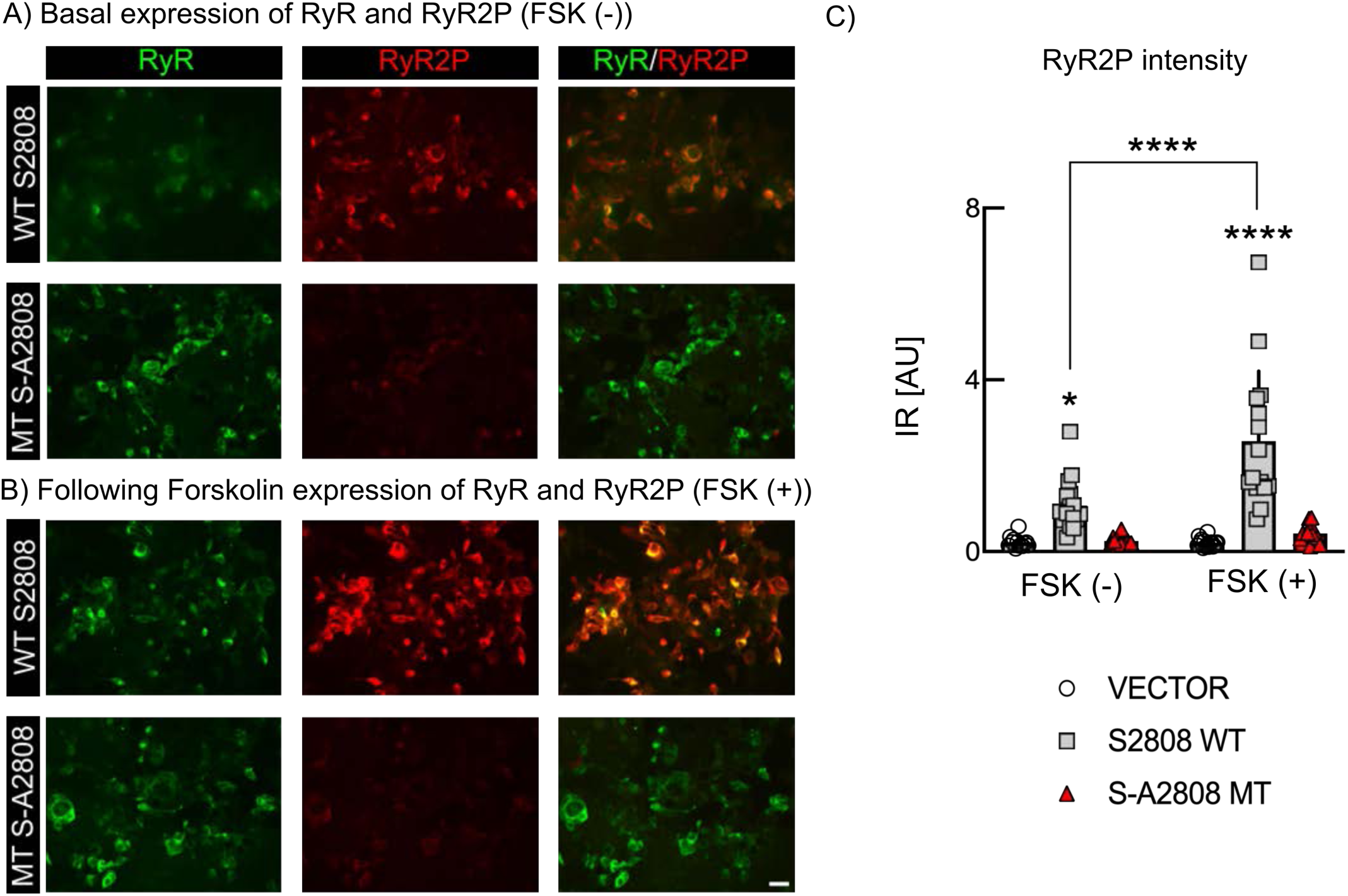
Validation of RyR2P antibody. Representative confocal photomicrographs of HEK cells under (**A**) basal conditions and (**B**) following forskolin treatment. Cells expressed either wild-type RyR2 (WTS2808) or a mutant construct where S2808 is converted to an alanine (MT S-A2808) to prevent phosphorylation. Forskolin treatment increased RyR (*green*) staining in WT 2808, but did not increase RyR2P (*red*) staining in MT S-A2808 cells. (**C**) Quantification of RyR2P IR in HEK cells (Two-way ANOVA, Interaction: F_(2,82)_=9.717, p=0.0002, treatment: F_(1,82)_=12.54, p=0.0007, transfection: F_(2,82)_=46.05, p<0.0001; followed by Tukey’s multiple comparison Test; FSK (-) Vector vs. FSK (-) WT S2808, p=0.0244; FSK (-) WT S2808 vs. FSK (-) MT S-A2808, p=0.0274; FSK (+) Vector vs. FSK (+) WT S2808, p<0.0001; FSK (+) WT S2808 vs. FSK (+) MT S-A2808, p<0.0001); FSK (-) WT S2808 vs. FSK (+) WT S2808, p<0.0001. Scale bar = 20 μm.

**Figure 3, supplemental figure 1.**
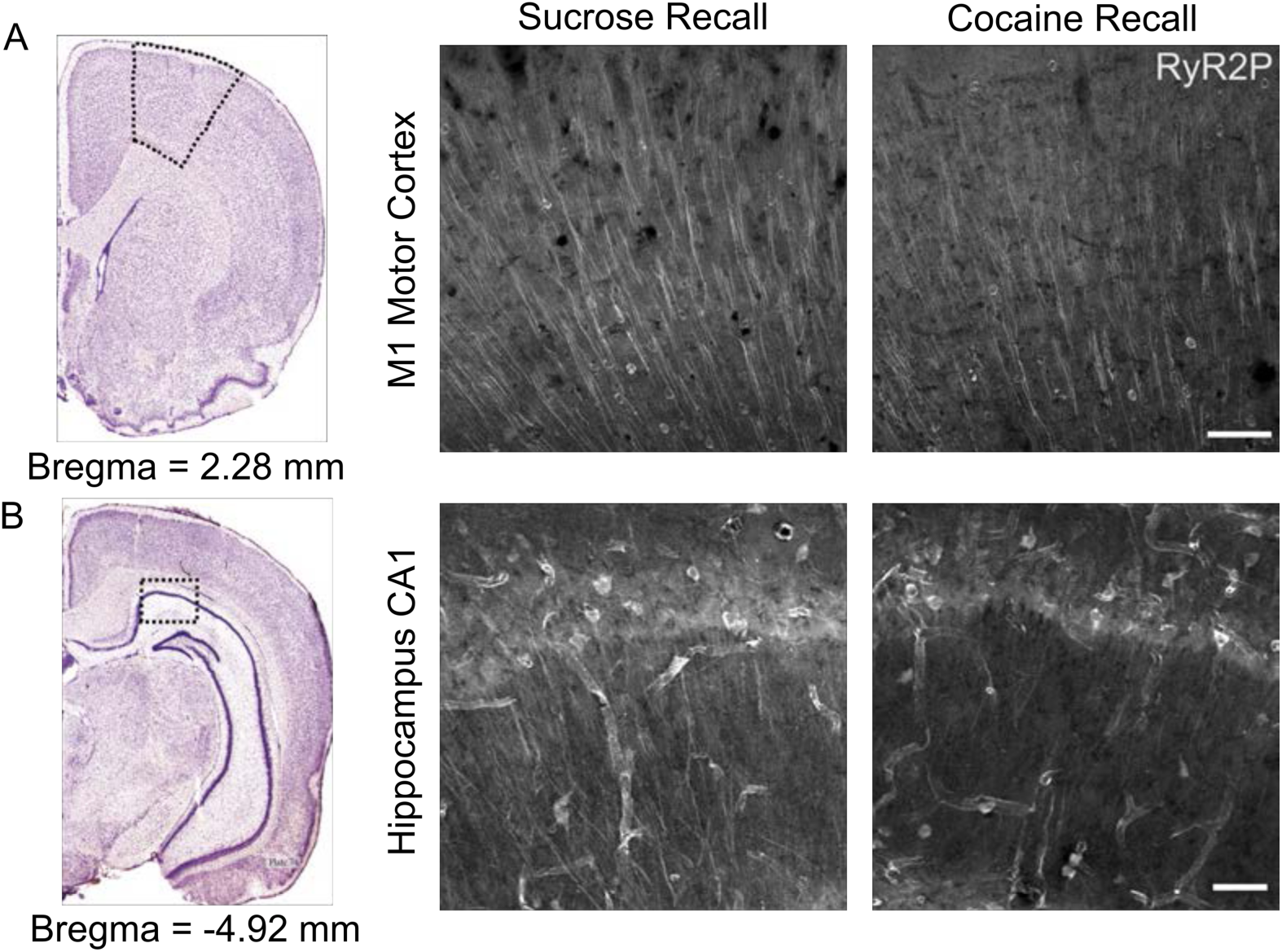
Spatial localization of RyR2P in M1 and CA1 pyramidal neurons following recall of natural and drug rewards. 20x confocal photomicrographs showing RyR2P (*white*) in (**A**) M1 motor cortex and (**B**) CA1 of the hippocampus of sucrose-SA and cocaine-SA rats that underwent recall testing. Scale bars: (A) = 100 μm; (B) 50 μm

**Figure 5, supplemental figure 1.**
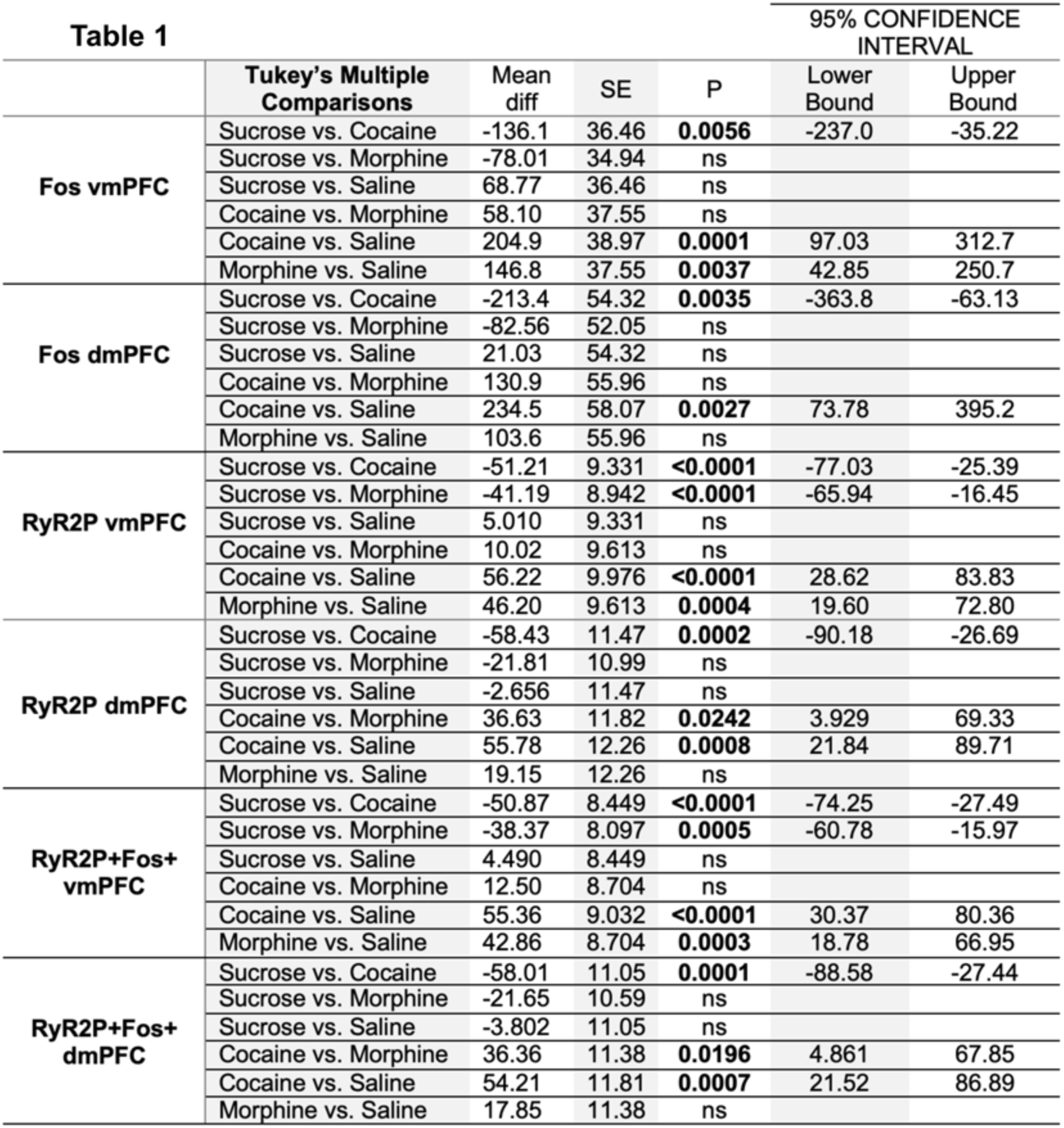
Table 1 shows the post hoc Tukey’s multiple comparison test from figure 5.

**Figure 5, supplemental figure 2.**
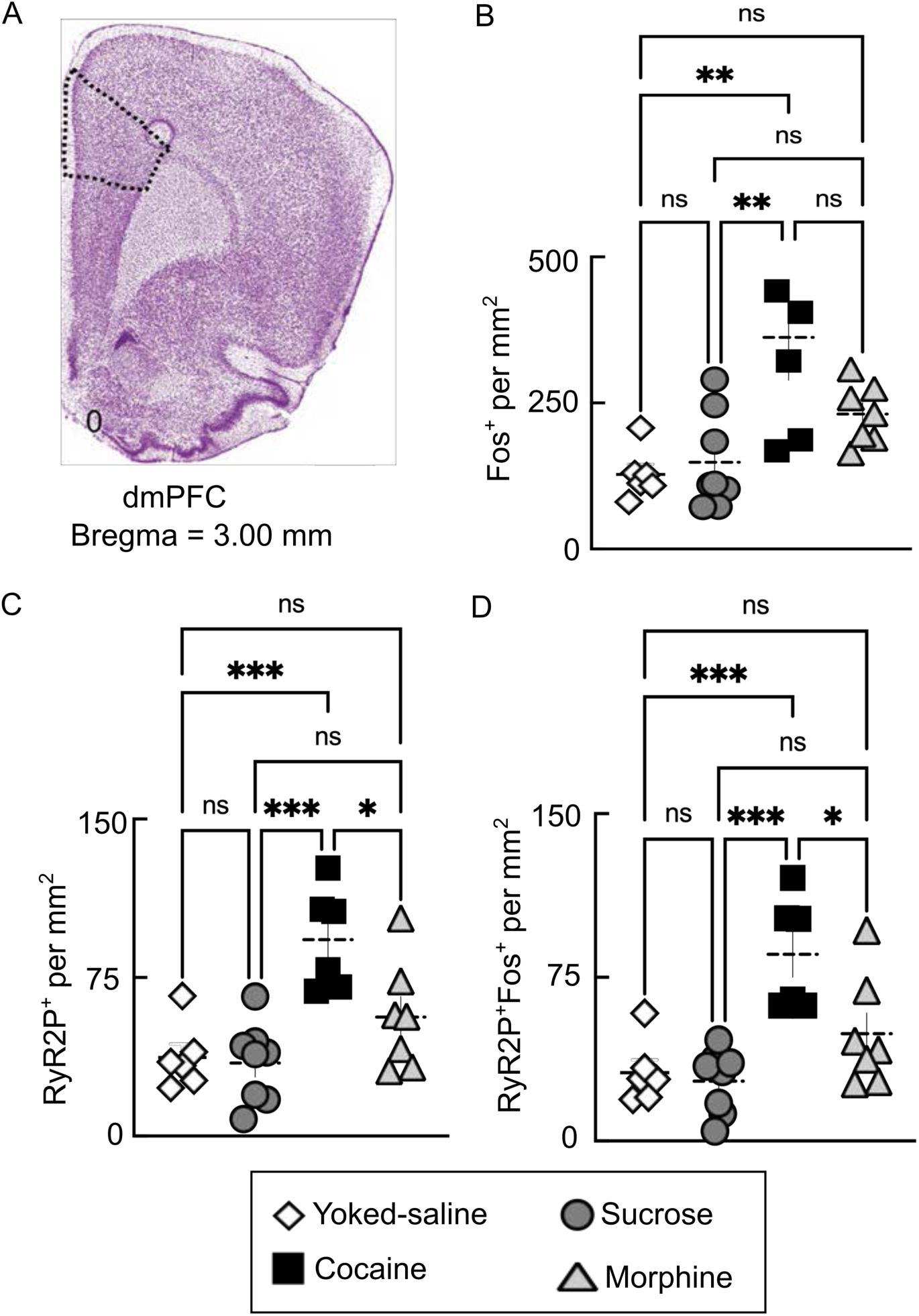
Cocaine and morphine recall related changes in dmPFC. (**A**) Atlas plate showing dmPFC (*black dotted outline*) at Bregma = 3.00mm. Graphs summarizing dmPFC differences in the expression of (**B**) Fos+ (F_(3,23)_=7.004, p=0.0016), (**C**) RyR2P+ cells (F_(3,23)_=10.25, p=0.0002), and (**D**) RyR2P+Fos+ cells (F_(3,23)_=10.71, p=0.0001). Note the enhanced numbers of RyR2P+Fos+ cells in cocaine (but not morphine) recall. One-way ANOVA followed by Tukey post hoc. *=p≤0.05, **=p≤0.01, ***= p≤0.001. N=6-8 rats per experimental group with 4 technical replicates per rat. Error bars represent S.E.M.

**Figure 5, supplemental figure 3.**
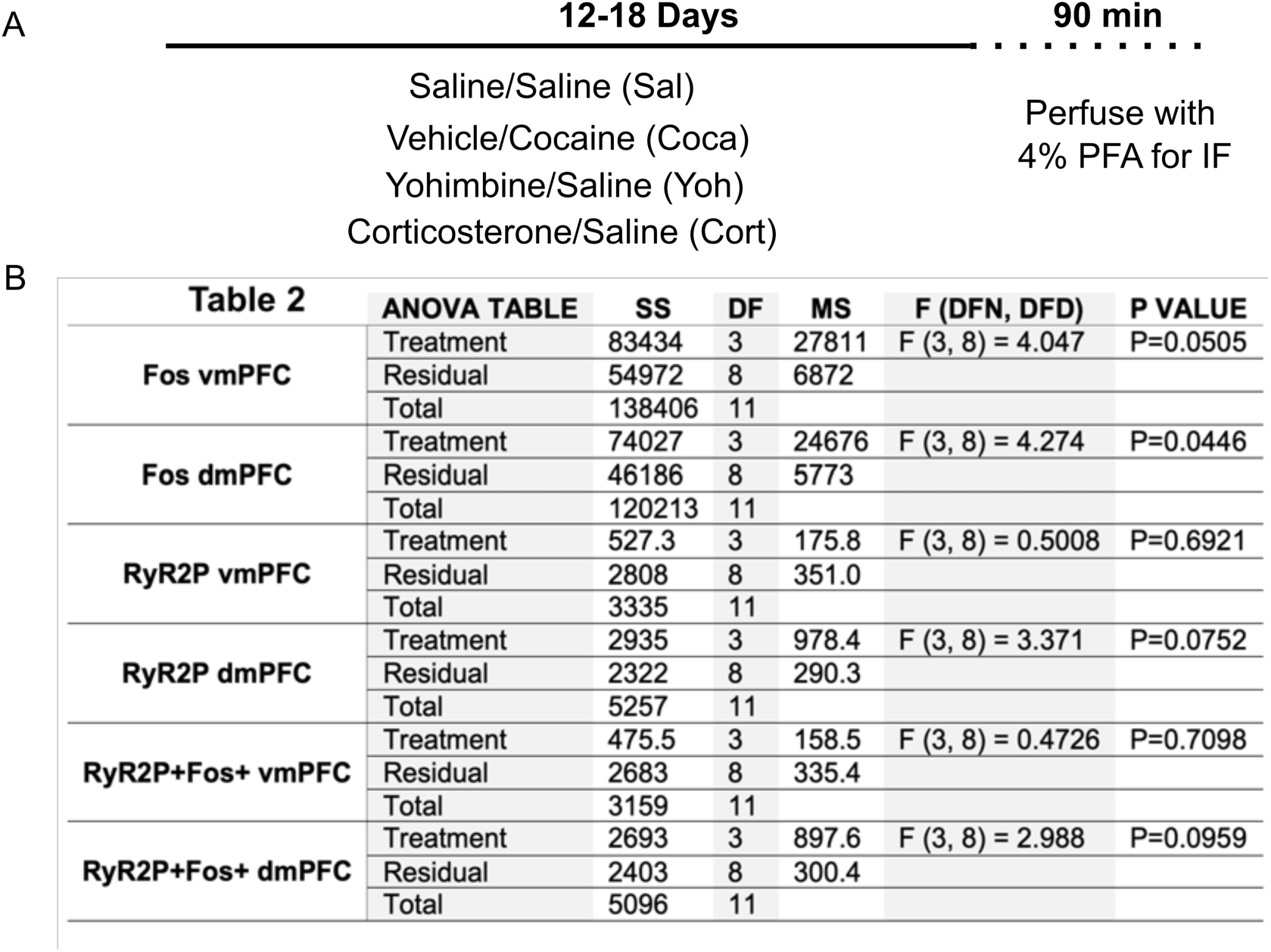
Non-contingent cocaine or stress does not increase RyR2P in the PFC. (**A**) Experimental workflow for chronic (12-18d) non-contingent (i.p. injections) treatments of: saline/cocaine (Coca; 20mg/kg), saline/corticosterone (Cort; 2mg/kg), saline/yohimbine (Yoh; 2mg/kg) or sterile saline (Sal) as a control. Following treatment, rats were perfused with 4% PFA to examine RyR2P expression in the vmPFC. (**B**) Table 2: summary ANOVA table for Fos1+, RyR2P+, and RyR2P+Fos+. N=3 rats per experimental group with 4 technical replicates per rat.

